# Life in sediments fosters ‘sexual’ speciation in *Shewanella baltica*

**DOI:** 10.64898/2026.03.14.705985

**Authors:** Víctor Fernández-Juárez, Francisco Salvà-Serra, Guillem Seguí, Alberto J. Martín-Rodríguez

## Abstract

Understanding how intra- and interspecific differentiation arises in natural microbial populations is central to explaining the processes that drive bacterial evolution. Motivated by the co-occurrence of several genospecies closely related to *Shewanella baltica* in Baltic Sea sediments, we investigated the genomic structure of this species complex across fine spatial scales. We analyzed 112 genome sequences from strains collected across multiple sediment cores and depths (0-6 cm) at Vaxön (Stockholm archipelago, Sweden), including sympatric isolates from this site as well as earlier isolates and allopatric strains from other locations in the Stockholm region obtained from both sediments and the water column. Using a reverse-ecology population genomics approach, we found that these strains form a species complex that resolves into three cohesive evolutionary lineages (G1, G2, and G3). Each lineage is characterized by extensive gene turnover, driven largely by horizontal gene transfer (HGT), and displays distinct genomic signatures of metabolic specialization. While G1 consists predominantly of a single species (*S. baltica*), G2 and G3 comprise a diverse set of divergent genospecies, many of which are repeatedly recovered from sediment samples. Patterns of homologous recombination indicate that speciation within G2 and G3 is primarily recombination-driven (‘sexual’), and that both groups derive from a common ancestor. Together, these results capture a snapshot of early-stage speciation within a shared ecosystem and provide insight into the mechanisms that diversify sympatric, recombining bacterial populations, with a sediment-associated lifestyle likely promoting this process.

## INTRODUCTION

Defining how microbial diversity is structured is central to understanding microbial evolution. Analyses of bacterial whole-genome sequences and metagenome-assembled genomes have consistently shown that microbes tend to form sequence-discrete clusters with high intra-group average nucleotide identity (ANI), typically ≥95%, corresponding approximately to 70% of DNA-DNA hybridization (DDH) values ^1^, and separated by gaps in the range of 83-95% ANI ^2^. While it remains an open question whether a continuum of genomic diversity among prokaryotes exists ^3^, these distinct clusters are broadly accepted as corresponding to what we refer to as ‘bacterial species’. Species circumscription thresholds are, however, somewhat blurry, with a frequently defined ‘gray zone’ around the 95-96% ANI window ^4^, and more stringent cutoffs (e.g., ANI ≥96% or taxon-specific thresholds) sometimes providing greater taxonomic coherence ^5, 6^.

A challenge for the definition of bacterial species relies on how bacterial populations evolve. As asexual organisms, bacteria reproduce clonally by binary fission. Over time, DNA mutations may arise during DNA replication and lead to clonal speciation. Nevertheless, this appears to be a rarity in nature, as less than 10% of all bacterial species have been estimated to be truly clonal ^7^. Instead, horizontal gene transfer (HGT) through various mechanisms has been consistently shown to represent, overall, a more powerful evolutionary driver ^8–10^. Albeit HGT can occur across distant phylogroups, including across kingdoms ^11, 12^, large-scale genetic exchange is rare between distant genomes ^13^. Thus, analogously to the fundamentals (DNA recombination) and limits (interbreeding) of sexual reproduction in eukaryotes, barriers in large-scale HGT across bacteria could be regarded as species boundaries, and speciation driven by HGT is often referred to as ‘sexual speciation’ ^14^. Indeed, the blurry ANI threshold delimitating species boundaries mentioned above has been proposed to emerge from the ANI limit (90-98%) in which gene flow is interrupted across distinct lineages ^7^. Evidence suggests that HGT goes hand in hand with ecological adaptation, with HGT being promoted in bacterial populations sharing the same habitat or lifestyle ^15–17^. This view is consistent with several speciation models that treat species as metapopulation lineages whose cohesion reflects both genetic exchange and shared ecological pressures, occupying different positions along a continuous speciation spectrum shaped by gene flow, recombination, and selection ^18–21^.

Renowned for their exceptional metabolic versatility, *Shewanella* species are ubiquitously distributed in water ecosystems and redox-stratified environments like aquatic sediments, where they play important roles in nutrient cycling and biogeochemistry ^22–24^. The Baltic Sea, a semi-enclosed and highly stratified water body with sharp oxygen gradients across the water column and a longitudinal salinity gradient, is home to *Shewanella baltica*, a species that thrives under brackish conditions. Different *S. baltica* ecotypes, defined as independent monophyletic clusters occupying specific ecological niches ^25, 26^, have been described across the oxygen gradient of the water column ^27^. These *S. baltica* populations undergo extensive homologous recombination, facilitating genetic exchange among strains, enhancing metabolic capabilities likely contributing to ecological fitness, and promoting strain diversification ^28^. Although the high rates of HGT in *S. baltica* have been postulated to make ‘sexual’ speciation possible ^27–31^, factual evidence of new species formation has remained elusive.

Our team has devoted efforts to the genomic characterization of *Shewanella* spp. from water and sediments of the Baltic Sea and other aquatic environments around Stockholm, which have led to the identification of several novel species closely related to *S. baltica*, i.e., *S. scandinavica*, *S. vaxholmensis*, and *S. septentrionalis*, collectively referred to as the ‘*S. baltica* complex’, with *S. hafniensis* occupying a borderline position ^32, 33^. Thus, intrigued by the co-occurrence of *S. baltica* and multiple intimately related novel genospecies in sediments, we set out to examine the genomic variation within this species complex. We examined the genomic content of arbitrarily selected strains from sympatric *Shewanella* populations sampled at different depths (0, 2, 4, and 6 cm) in sediment cores collected at Vaxön in the Stockholm archipelago. We also included strains obtained from earlier sampling at the same site and arbitrarily selected allopatric strains from other locations in the Stockholm region, resulting in a total dataset of 112 genome sequences. Using a comprehensive suite of state-of-the-art bioinformatic inferences, we delved into the genomic structure of the *S. baltica* complex and the evolutionary forces shaping and maintaining the observed phylogenetic and ecogenomic patterns. The results of these analyses led us to propose that *S. baltica* genomic divergence into separate taxa is largely ‘sexual’, is stimulated by a sediment-associated lifestyle, and involves the emergence of coexisting, stable strain lineages that are consistent with the standards for the designation of bacterial species ^34^.

## RESULTS

### Surficial sediments in Vaxön are taxonomically and functionally homogenous

Our study began with the collection of four surficial sediment cores, termed SP0, SP1, SP2, and SP4 (∼7-8 cm deep) from the shoreline of Vaxön, an island in the Stockholm archipelago (12 August 2023; Fig. S1). Environmental DNA was extracted from sediments at 0, 2, 4, and 6 cm depths for metagenomic analysis. Across these depths, metagenomics revealed no significant changes in bacterial community composition or functional pathways (Fig. S2A-E; p > 0.05). *Actinobacteriota* dominated the communities (18.1-38.5% of reads), followed by *Pseudomonadota* (17.0-25.0%), *Bacteroidota* (1.3-2.4%), and *Planctomycetota* (1.0-2.2%) (Fig. S2A). Neither alpha- nor beta-diversity, assessed using Shannon indices and Euclidean distances, respectively, differed significantly with depth (p > 0.05; Fig. S2A, B). Functional annotation of metagenomic reads based on KEGG pathways showed no depth-dependent clustering (p > 0.05, Kruskal-Wallis test; Fig. S2C, D). Predominant pathways (log_10_ GPM > 7) included ABC transporters, quorum sensing, DNA replication and repair, glycolysis/gluconeogenesis, fatty acid metabolism, and purine metabolism (Fig. S2E). *Shewanella* spp. populations accounted for 0.04-0.21% of total bacterial reads (Table S1), representing more than 2.3% of *Gammaproteobacteria* and ranking among the 100-300 most abundant genera detected (out of >13,000). However, *Shewanella* abundance did not differ significantly across sediment depths (Fig. S2F; p > 0.05). Together, these data indicate that (i) bacterial community composition remains largely unaltered across the upper 6 cm of sediments, and (ii) *Shewanella* is not a particularly dominant taxon in the surficial sediment community.

### A sympatric lifestyle within surficial sediments fosters genotypic divergence in *Shewanella* populations

From the same sediment samples, we retrieved 60 randomly selected sympatric *Shewanella* strains (designated with the prefix ‘VAX’; Table S2). We complemented this collection with 18 *Shewanella* strains obtained during previous sampling and isolation efforts in the same area in 2021 (prefix ‘SP’), as well as randomly selected allopatric strains from sediment (n = 19) and water (n = 15) samples collected from distinct locations in and around Stockholm between 2021 and 2023. These sites included Nynäshamn (n = 11; prefix ‘N’), Tyresö (n = 12; prefix ‘T’), Lidingö (n = 8; prefix ‘L’), and Hagaparken (n = 3; prefix ‘H’) (Fig. S1). In total, we analyzed 112 *Shewanella* genomes, including previously sequenced strains ^32, 33^, of which 32 were complete genomes (all of which belonged to representative members of the *S. baltica* complex) (Table S2).

To assess the preliminary taxonomic diversity of the *Shewanella* isolates retrieved from Vaxön, comprising 78 isolates from the 2021 and 2023 sampling occasions combined, whole-genome sequences were submitted to the Type Strain Genome Server (TYGS) ^35^, which uses 70% digital DNA-DNA hybridization (dDDH) as threshold for species designation. Based on this indicator, 47% of the isolates were affiliated with *S. baltica* (n= 37), 10% to *S. vaxholmensis* (n = 8), 5% to *S. scandinavica* (n = 4), and 1% to *S. septentrionalis* (n = 1)*, whereas a remarkable 32% could not be* confidently assigned to any known species *(n = 25)*, but exhibited the closest similarity to *S. baltica* (Fig. 1A). Thus, ∼95% of the Vaxön isolates in this dataset were taxonomically very closely related, with the remaining strains affiliated with *S. hafniensis*, *S. oncorhynchi*, and *S. holmiensis* (Fig. 1A). Pairwise ANIb comparisons between all these strains exhibited values ranged from 89.6% to 100%, with two clear ANI gaps observed between the interactions *S. oncorhynchi* vs. *S. hafniensis* (91.0-93.2%) and *S. hafniensis* vs. *S. baltica* complex members (94-94.5%) (Fig. 1A). Of note, we found a distinct pattern within the *S. baltica* complex characterized by a continuous bimodal distribution with 60% (3,743 pairwise comparisons) falling within the 94.5%-95.9% range, while 28% (1,748 comparisons) clustered between 95.9%-97.2% (Fig. 1A).

**Figure 1.**
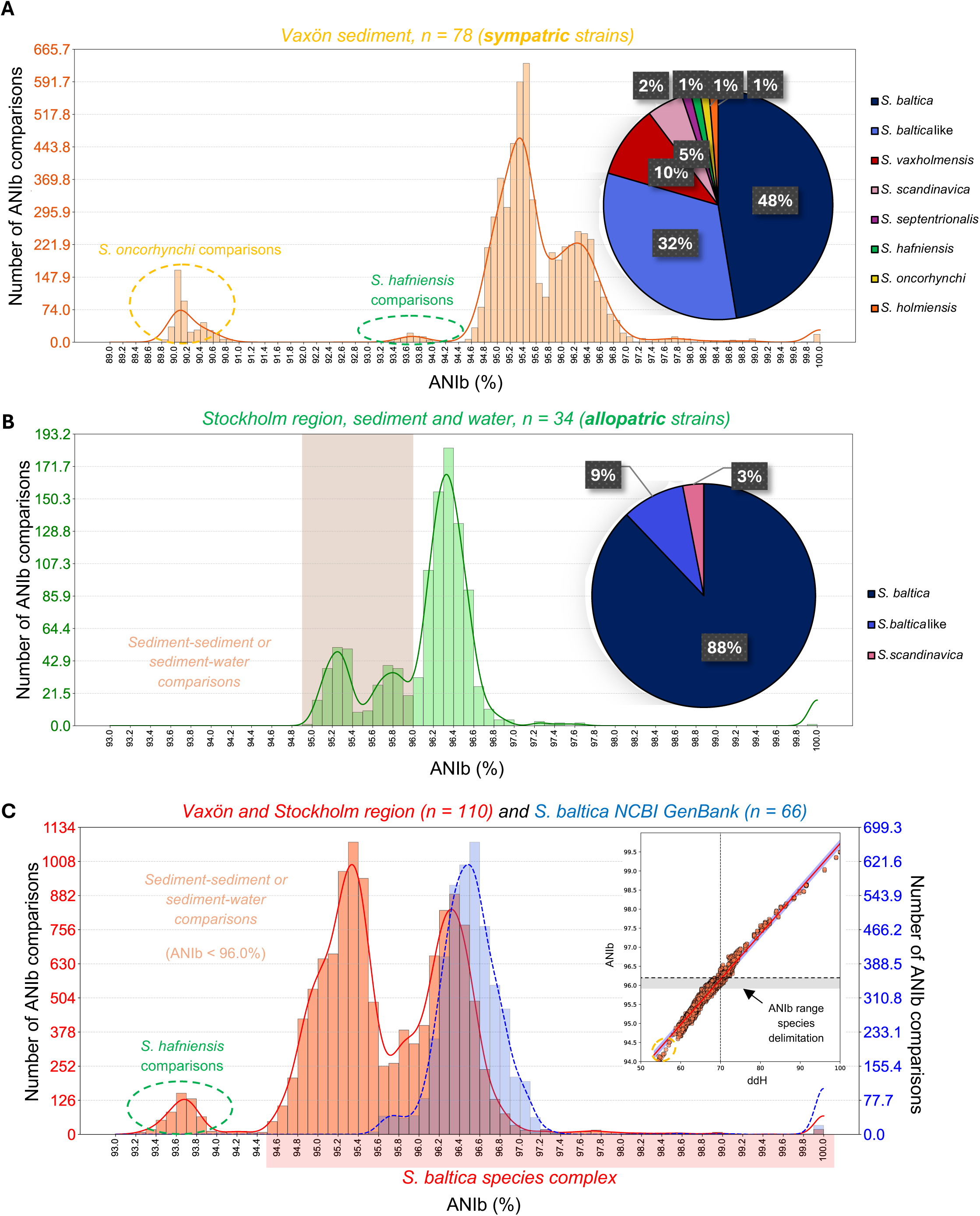
Genomic diversity of *S. baltica* complex genomes based on ANIb relatedness. (**A-C**) Frequency distributions of pairwise ANIb comparisons (0.1% windows) for (**A**) strains recovered from Vaxön sediment cores in 2023 (n = 60) and from previous sampling occasions in the same region (n =18); (**B**) allopatric strains retrieved from other regions in or around Stockholm city: sediments (n = 19) and water (n = 15); and (**C**) the entire dataset, including both our isolates (108 *S. baltica*-like and 1 *S. hafniensis*, excluding *S. oncorhychi*) along the type strain *S. baltica* CECT 323^T^(in red), and those available in the NCBI GenBank (n = 66, shown individually in blue for comparison). In (**A**), color circles denote the interactions with the closest species to the *S. baltica* complex recovered from our study, i.e., *S. oncorhychi* (n = 3) and *S. hafniensis* (n = 1). In (**A-B**), pie charts showing the affiliation of the isolates recovered in the as determined by TYGS were added as insets. In (**B),** the brown frame highlights the interaction with closely related strains from the sediment. In (**C**), the all-vs-all genome-to-genome distance calculator (GGDC) analysis is embedded within the ANIb comparison, showing that ANIb values corresponding to the 70% digital DNA–DNA hybridization (dDDH) species threshold fall within 95.8–96.1%, thereby defining species boundaries within the species complex.

We extended our analysis to a broader set of 34 allopatric *S. baltica* complex strains isolated from sediment and water samples collected at distinct locations in Stockholm and its surrounding areas (Fig. S1). In this set, 88% of the strains (n = 29) could be unambiguously affiliated to *S. baltica* by TYGS, whereas 9% (n = 3) could not be affiliated to a known species, and 3% (n = 1) circumscribed within *S. scandinavica* (Fig. 1B). Here, the ANIb distribution revealed a trimodal pattern with 15.4% of interactions (168 pairwise comparisons) falling within the 94.8%-95.5% range, 12.4% (135 pairwise comparisons) within 95.5%-96.0%, and 68.2% (743 pairwise comparisons) within 96.0%-97.0% (Fig. 1B). The first two ANI peaks reflected interactions between sediment-sediment or sediment-water genome pairs, whereas the highest density peak (96.0%-97.0%) was observed exclusively among water-water genome comparisons, suggesting a potential role of sediments in driving genomic diversification.

ANIb pairwise comparisons across all genomes, integrating the local Vaxön dataset (sympatric) with the Stockholm regional dataset (allopatric), support the bimodal distribution previously mentioned, showing a high average shared genome fraction (84.8 ± 3.63%; Fig. S3A). The lower peak (94.5-95.6 ANIb%) accounted for ∼50% of comparisons (n = 6,383), mostly involving sediment-derived isolates, with the sole exception of *S. septentrionalis*, isolated from surficial shore water near the sediment layer ^33^. Notably, no NCBI GenBank *Shewanella* genome sequences fell within this range (reviewed on 31 May 2025, Table S3), highlighting the novelty of this population. The upper peak (centered ∼96.3 ANIb%) encompassed 32% of comparisons (n = 4,148), predominantly *S. baltica* strains, closely overlapping publicly available *S. baltica* genomes (∼96.5 ANIb%; Fig. 1C). This range, bordering the canonical ∼96% species delineation threshold, was contextualized by FastANI-based intra-specific comparisons within other well-represented *Shewanella* clades (*S. algae*, 219 genomes; *S. xiamenensis*, 98 genomes; *S. oncorhynchi*, 32 genomes; Table S3), which displayed unimodal ANI distributions with narrower dispersion and consistently higher intra-specific similarity (>97-98% ANI). In contrast, *S. baltica* spanned a broader ANI range extending toward the species boundary zone. Together, this shows the exceptional intra-specific diversity of *S. baltica* (Fig. S3B). Corresponding ANIb values at the 70% dDDH species threshold across the *S. baltica* complex fell within 95.8-96.2% (Fig. 1C, inset), defining species boundaries within this complex. Therefore, more than half of ANIb values fell within 94.5-96.2%, a transition zone consistent with the typical ANI gap for species delineation ^4^, and thus, representing a possible case of ongoing sympatric speciation in sediments.

### Multiple novel genospecies coexisting in surficial sediments provide a snapshot on *Shewanella* speciation

To assess the extent of speciation within our dataset, we performed pairwise ANIb and dDDH comparisons and reconstructed phylogenies based on the core genome and core proteome content. Thus, genomic species (genospecies) were defined based on current standards ^34^, using the following criteria: i) ANIb ≥ 95.8-96.2% (all vs. all); ii) dDDH ≥ 70% (all vs. all); and iii) phylogenomic coherence between core genome and core proteome trees. All-against-all ANIb comparisons defined three main groups (G), namely, G1, comprising mostly *S. baltica* strains plus several unassigned strains; G2, closely related to *S. scandinavica*; and G3, related to *S. vaxholmensis* and *S. septentrionalis* (Fig. 2A). ANIb values between groups showed clear boundaries supporting this clustering (<96%), with intergroup similarities ranging from 94.89-95.97% (G1–G2), 94.50-95.91% (G1–G3), and 95.10-95.71% (G2–G3) (Fig. 2A). Genome-wide differentiation within the *S. baltica* complex was assessed using SNP-based sliding window analyses, and pairwise p-distances ranged from 0 to 0.5 (Fig. 2B). Values above 0.25, observed within and between groups, indicate high differentiation, whereas values near 0, more frequent in G2 and G3, indicate emerging species clusters (Fig. 2B), aligning with the ANIb group classification.

**Figure 2.**
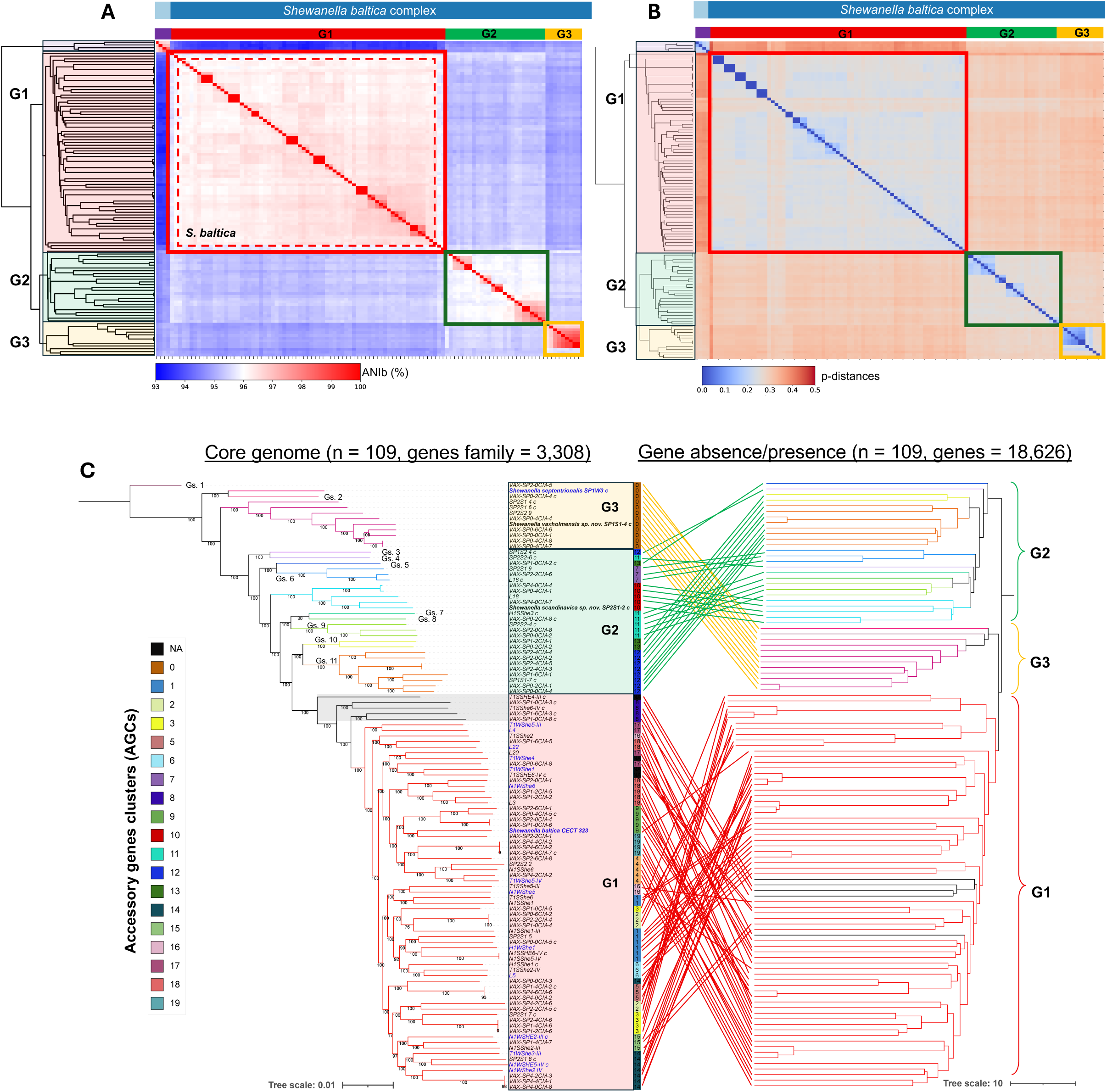
ANI clustering, genome-wide, and pangenome analysis of the *S. baltica* complex from the Stockholm region. (**A**) Heatmap of all-vs-all ANIb values were calculated for the *S. baltica* complex. The dashed red frame within G1 denote the strains affiliated with *S. baltica*. (**B**) Heatmap showing pairwise genome-wide differentiation between strains, based on SNPs extracted from whole-genome alignments. SNPs were grouped into overlapping 10 kb sliding windows with a 2.5 kb step size. For each window, the proportion of SNPs with discordant genotypes (p-distance) was calculated for all genome pairs, producing normalized values from 0 (identical) to 0.5 according to the genomic divergence. In (**A**) and (**B**), as the closest relative, *S. hafniensis* (SP1S1-9 and 3 additional from NCBI, Table S3) was added as an outgroup represented in purple. (**C**) Combined core genome and accessory genome tree constructed using a 90% identity and 90% coverage threshold based 3,310 core gene and protein families from the 109 genomes (including *S. baltica* CECT 323^T^ as reference). For building the accessory tree, pairwise Euclidean distances between isolates were calculated, and an average-linkage (UPGMA) method was applied. Blue labels indicate genomes isolated from the water column. Within G2 and G3, branch colors indicate distinct species previously described: *S. scandinavica*, *S. vaxholmiensis*, and *S. septentrionalis*, as well as putative genospecies (“Gs.”). Species and genospecies were delineated as explain in the main text. Accessory gene groups (AGCs) detected by Mandrake are indicated in the tree, each marked by distinct numbers and colors. Lines link corresponding genomes between the core genome and accessory genome trees, illustrating their topological relationships. Small numbers on the branches denote bootstrap support values after 1,000 replications. In all the plots, groups are differentiated by color, i.e., red for G1, green for G2, and yellow for G3. The grey shadow denotes the genomes could not be affiliated to *S. baltica*.

Our phylogenomic reconstructions, based on both core genome and core proteome content, comprising 3,308 core family genes (3,399,949 bp) or proteins (1,131,112 aa) from a pangenome of 18,626 genes (representing ∼75% of the core genes in each genome), were highly congruent with one another. Their topology was consistent with inferences drawn from overall genome relatedness indexes (OGRIs) previously introduced (Fig. 2C, S4A). In total, we identified 11 putative novel genospecies within groups G2 and G3, specifically, 9 within G2 and 2 within G3 (Fig. 2C), in addition to the previously proposed species *S. septentrionalis*, *S. vaxholmiensis*, and *S. scandinavica* ^32, 33^. Several genome sequences from different time points (2021 - ‘SP’ prefix, and 2023 - ‘VAX’ prefix) were affiliated with the same species or genospecies, e.g., various *S. vaxholmensis* strains (G3) and diverse isolates belonging to a novel putative genospecies within G2, e.g., Gs. 11 (Fig. 2C). This finding evidenced stability over time of divergent lineages in this sediment ecosystem. In contrast, no evident grouping of strains was observed by geography or sediment depth, the former indicating widespread distribution of lineages in the studied region (Fig. 2C), and the latter being consistent with the apparent taxonomic and functional homogeneity inferred from our metagenomic analyses. Thus, our phylogenomic analyses show that sympatric speciation by members of the *S. baltica* complex might be driven by the close coexistence of strains in sediments, which could be promoted by extensive lateral gene exchange in this habitat.

### High rates of lineage-specific gene turnover drive accessory genome diversification in the *S. baltica* complex

To contextualize gene exchange in the *S. baltica* complex, we assessed the distribution of accessory genes across the phylogeny. We integrated an accessory genome tree derived from gene presence/absence profiles with the core genome phylogeny, which revealed distinct and group-specific accessory gene clusters (AGCs) which, overall, were congruent with the topology of the phylogenomic reconstructions (Fig. 2C). Species within G3 shared a highly similar AGC, whereas nearly every species or genospecies within G2 exhibited a distinct AGC from one another. In contrast, G1 strains displayed a much higher diversity of AGCs, up to 16, which did not consistently align with the core genome tree topology (Fig. 2C). The high number of AGCs observed in the *S. baltica* complex is consistent with its evidently open pangenome, encompassing up to 87% of the total pangenome and including over 15,000 accessory genes (Fig. 3A), constituting ∼25% of each genome. Despite G1 forming a cohesive cluster primarily composed of a single species, *S. baltica*, it exhibited substantially higher accumulated accessory genome content than G2 and G3 combined (Fig. 3A), exceeding that observed in other *Shewanella* species such as *S. algae* and *S. xiamenensis*, and comparable to that of *S. oncorhynchi* (Fig. S4B), indicating an elevated rate of gene turnover in G1 relative to G2 and G3, which display a less open pangenome (Fig. 3A).

**Figure 3.**
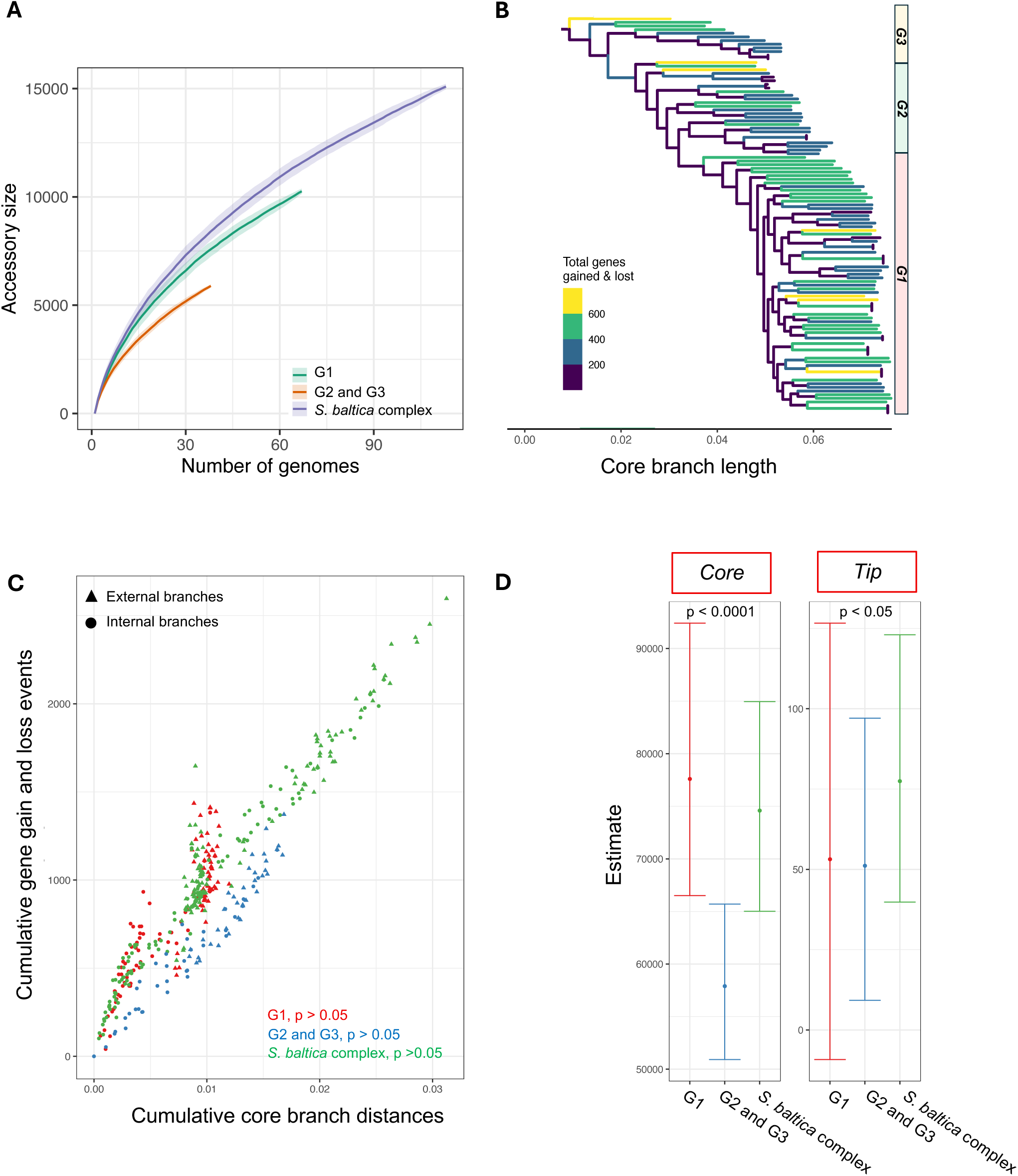
Gene gain, loss, and exchange patterns in the *S. baltica* complex pangenome. (**A**) Accumulation curves of accessory genome content of the members of the *S. baltica* complex from this study (n = 109) and dissected by evolutionary group. (**B**) Core genome tree, with branches colored according to the number of gene gain and loss events inferred using maximum parsimony. (**C**) Cumulative number of gene gain/loss events plotted against cumulative branch length, and (**D**) gene exchange parameters estimated by Panstripe. “Core” reflects the association between gene gain/loss events and core phylogeny branch length, and “tip” indicates gene exchange events occurring specifically at terminal branches. p-value reflects the significance of turnover events associated with the core and the tip within the groups compared. In (**A**), (**C**), and (**D**), to minimize potential biases in comparisons, we merged G2 and G3 (n = 38), which, although evolutionary distinct, displayed significantly lower accessory content and rates of gene turnover compared to G1.

We next studied the gene turnover in *S. baltica* complex using the Panstripe pipeline ^36^. Overall, the *S. baltica* complex exhibits 200-800 gene gain/loss events per genome (Fig. 3B), with G1 showing significantly more events (352 ± 124) than G2 and G3 (250 ± 168; p < 0.05; Fig. 3B), consistent with its broader accessory gene content (Figs. 2C, 3A). Older (internal) and newer (external) lineages displayed similar rates of gene turnover along their corresponding phylogenetic branches (p > 0.5; Fig. 3C), yet a pronounced ‘tip effect’ was detected across the complex, with turnover events concentrated on terminal branches, indicating that recently diverged lineages experience elevated levels of gene turnover between them (p < 0.05; Fig. 3D). Gene turnover was strongly associated with core genome divergence of G1 relative to G2/G3 (p < 0.001; Fig. 3D), showing that recently evolved, genetically distinct lineages significantly contribute to overall gene turnover. Although the exceptionally high gene turnover in the *S. baltica* complex, particularly in G1, does not fully explain the emergence of new genospecies in G2 and G3, it evokes the notion of ecological differentiation.

### Metabolic specialization mediated by HGT differentiates groups within the *S. baltica* complex

To investigate this potential functional differentiation, we first examined the overall distribution of genome functions across *S. baltica* complex strains using RAST-SEED annotation, assessing the proportion of annotated protein-coding genes assigned to each functional category. The resulting PCA of the functional profiles revealed three distinct and well-separated clusters, indicating clear functional differentiation among groups (PERMANOVA, pseudo-F = 12.82, p = 0.001; Fig. 4A). Second, to explore the genomic basis of this functional divergence, we quantified regions of genome plasticity (RGPs) to assess their contribution. Approximately 40% of accessory genes were located within putative RGPs, with each genome containing 140-380 putative HGT-derived genes (Fig. 4B). While no significant differences were observed in the number of RGPs or the taxonomic origin of HGT-derived genes among groups (Fig. S5A; p > 0.05), most originated from the family *Shewanellaceae* (58.7–75.6%), followed by diverse aquatic bacterial taxa including *Vibrionales*, *Pseudoalteromonadaceae*, *Alteromonadaceae*, and *Oceanospirillales* (Fig. S5B). G1 contained significantly more HGT-derived genes (328 ± 65 per genome) than G2 and G3 (250 ± 65 per genome; p < 0.05; Fig. 4B), consistent with its more open pangenome. Our PCA analysis based on COG annotations of HGT-derived genes revealed clear functional separation among groups, with G2 and G3 showing closer yet significantly distinct HGT profiles (PERMANOVA, pseudo-F = 9.65, p = 0.001; Fig. 4C), indicating group-specific gene acquisition that might drive ecological specialization.

**Figure 4.**
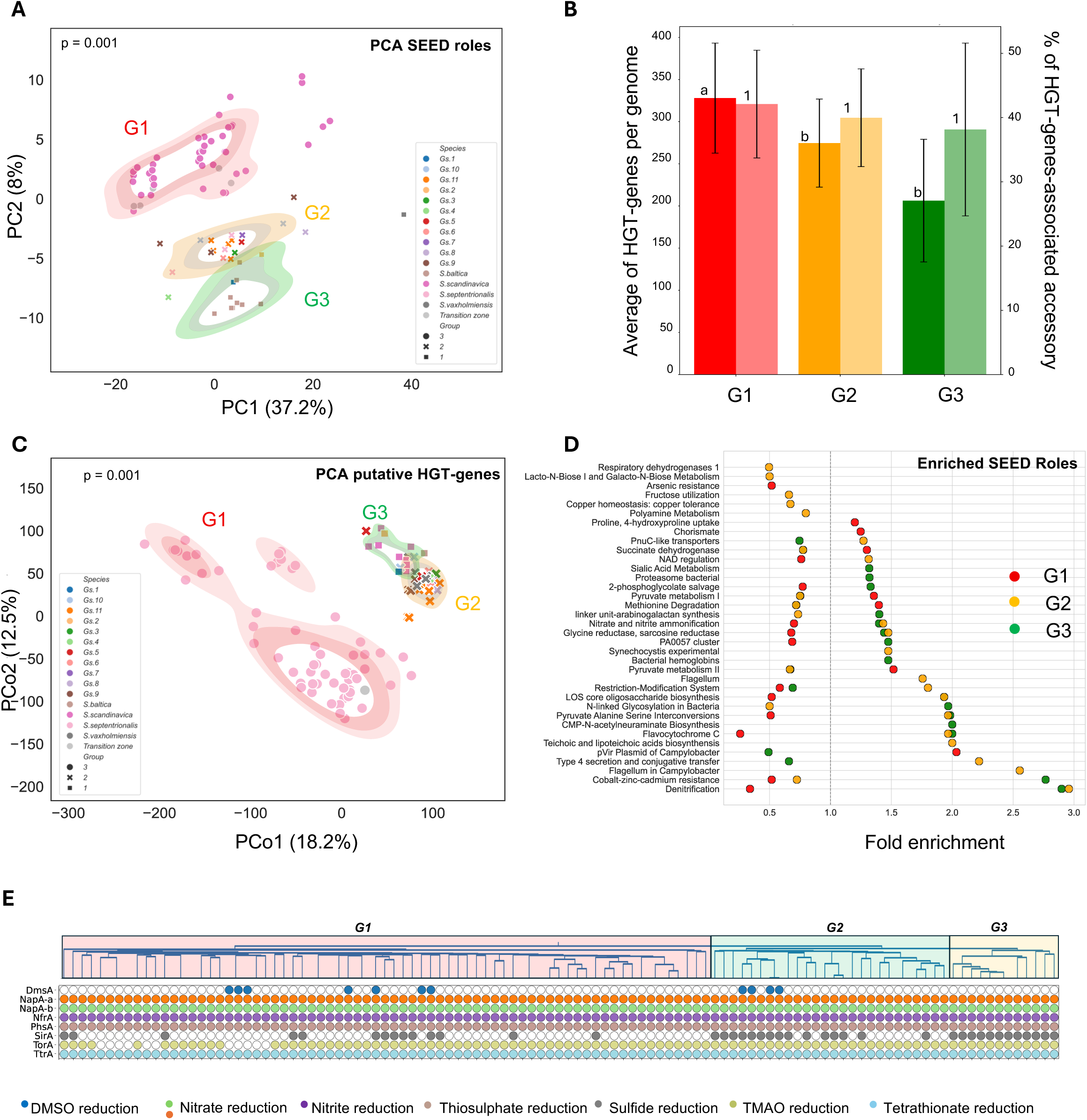
Horizontal gene transfer and its role in the ecological specialization of the *S. baltica* complex. (**A**) Principal component analysis (PCA) of functional profiles derived from SEED-annotated gene content of genomes. The analysis is based on the percentage of proteins associated with each functional category. (**B**) Number of HGT-genes per genome (first column) and the percentage of these genes (second column) over the total accessory of each genome across the described groups. Different letters or numbers denote if there are significant differences between the groups. (**C**) Principal coordination analysis (PCoA) of the COG-annotated genes located within the RGP, i.e., putative HGT-genes, across genomes of *S. baltica* complex. In (A) and (C), the first two principal components (PC1 and PC2) are shown, with the percentage of variance explained indicated on each axis, and PERMANOVA was conducted to assess group-level differences. Each point represents a genome, colored by species and shaped by group, with density ellipses highlighting group-specific clustering. (**D**) Functional enrichment analysis of SEED-annotated gene content across genomes. The x-axis shows fold change of the median value of that function per genome within each group relative to all other genomes, while the y-axis lists significantly enriched functions. For detecting the significant enriched SEED-annotated, respectively, Kruskal–Wallis tests were applied, with multiple testing corrected using the Benjamini–Hochberg false discovery rate (FDR). The specific enriched SEED-annotated and COGs by group are listed in Table S4 and S5. E) Presence/absence of reductases across *S. baltica* complex genomes. The plot depicts the distribution of reductases, with colored circles indicating presence and white cells indicating absence. Genes were grouped and color-coded according to which group of reductases they were affiliated by these functional categories as detailed in the legend. Genomes along the x-axis are ordered according to their relatedness in the ANIb.

With the aim to gain deeper insight into the acquired metabolic traits, we conducted a functional enrichment analysis to identify SEED functions and COGs significantly varying across groups and previously identified AGCs (FDR-adjusted p < 0.05; Fig. 4D; Tables S4, S5). *In silico* analyses indicated that G1 putatively possesses enhanced metabolic versatility in the degradation of diverse carbohydrate substrates relative to G2 and G3, with notable enrichment in pyruvate metabolism, methionine degradation, and succinate dehydrogenase activity (FDR-adjusted p < 0.05; Fig. 4D; Tables S4, S5). This was further supported by enrichment of L-arabinose utilization pathways in AGCs 4, 5, 6, 8, 9, 14, 15, 16, 17, and 18; lactose and galactose uptake in AGCs 4, 6, 8, 17, and 18, and trehalose utilization in AGCs 4, 9, 14, 15, and 18 (FDR-adjusted p < 0.05; Tables S4, S5). In contrast, G2 and G3 were enriched in denitrification and nitrogen metabolism functions (Fig. 4D). Additional enrichments included flavocytochrome C in AGCs 0, 10, 11, 12, and 13; pyruvate–alanine–serine interconversions in AGCs 0, 7, 10, 11, 12, and 13; and lacto-N-biose metabolism in AGC 7 (FDR-adjusted p < 0.05; Tables S4, S5), potentially linked to energy generation from endogenous carbohydrates and redox metabolism. Both groups shared enrichment in biosynthetic pathways for cell surface structures, such as lipo-oligosaccharide core oligosaccharides (Fig. 4D; Tables S4, S5). Further differentiation was observed, with G2 enriched in genes related to heavy metal resistance and restriction-modification systems, and G3 showing >2-fold enrichment in flagellar biosynthesis and type IV secretion systems (Fig. 4D; Tables S4, S5). KEGG pathway analysis assessing metabolic route completeness revealed no overall differences among groups, except for galactose biosynthesis and degradation pathways, which were more complete in G1 members (Fig. S6). Thus, although strains from G1, G2, and G3 were isolated from all sediment depths and metagenomic analyses showed no evidence of significant variations in microbial community composition or biological functions across the upper 6 cm of sediment, each group within the *S. baltica* complex exhibited distinct genomic signatures consistent with potential metabolic specialization, and reflecting diverse ecological strategies within the shared habitat.

Within this metabolic differentiation, we next focused specifically on energy metabolism, examining genes encoding reductases for alternative electron acceptors (AEAs) to evaluate potential differences in respiratory capacity. BLASTp searches were performed using reference *S. baltica* complex reductase catalytic subunits for nitrate (*napA*), nitrite (*nrfA*), thiosulfate (*phsA*), sulfite (*sirA*), tetrathionate (*ttrA*), dimethyl sulfoxide (DMSO; *dmsA*), and trimethylamine *N*-oxide (TMAO; *torA*) against all sequenced genomes (Fig. 4E, see Materials and Methods). Genes for nitrate, nitrite, tetrathionate, and thiosulfate reduction were conserved across all genomes (Fig. 4E). The *sirA* gene, encoding a sulfite reductase, was conserved in G3 but absent in roughly 50% of G2 and, particularly, among G1 members. We also observed loss of TMAO reductase genes in distinct G1 subspecies, alongside acquisition of DMSO reduction genes in multiple G1 and G2 strains (Fig. 4E; Table S6). DMSO reduction in *S. scandinavica* appears to be an acquired trait, as most strains (∼80%) encode a putatively functional DMSO reductase sharing 93.2% identity with that of *Shewanella colwelliana* (accession: WP_028764601). This enzyme is distinct from the *dmsA* variants in G1, which share only ∼30% amino acid identity, with the closest match to *Shewanella sediminis* (accession: WP_012143260). In both cases, *dmsA* and *sirA* orthologs were located within putative RGPs, highlighting the role of these regions in the acquisition or loss of energy metabolism-related traits. The gain and loss of reductase genes located in these regions might therefore promote metabolic differentiation among sympatric strains, supporting ecological specialization in a shared ecosystem.

Other mobile genetic elements, such as plasmids, might mediate important ecological adaptations and even speciation in different bacterial groups, such as *Rhizobium* ^37^. Here, we recovered 21 complete circular plasmid sequences from 32 closed genomes across the *S. baltica* complex, ranging from 2,677 to 120,678 bp. Plasmids were present in 15 of the 32 complete genomes, with each genome carrying 0-4 plasmids (Table S7). The identification of closely related plasmids in strains retrieved in 2021 and 2023 demonstrates plasmid stability in the sediment ecosystem (Fig. S7A). Although a small proportion of plasmids formed clusters based on FastANI relatedness, these clusters included representatives from all strain groups (Fig. S7A). COG annotation indicated enrichment in replication, mobilome, transcription, vesicular transport, and defense functions (Fig. S7B), but most plasmid genes remain uncharacterized (Table S7). Overall, while plasmid-borne genes can provide a range of yet-to-be-determined metabolic capabilities for their hosts, they are unlikely to be the main drivers of accessory genome differences in the *S. baltica* complex, with horizontally acquired chromosomal genes representing the primary source of genomic functional diversification.

### Recent genome exchange, coupled with sustained mutation rates, drives divergence within the ***S. baltica*** complex

While high gene turnover via HGT likely contributes to metabolic specialization in the *S. baltica* complex, additional evolutionary processes might also shape genomic diversification. To explore alternative drivers, we investigated whether recent genome exchange through homologous recombination plays a primary role, focusing on the *S. baltica* strains retrieved exclusively from sediment environments (n= 94). For that, we applied a recently described recombination detection method comparing the number of identical genes (>99.8% nucleotide sequence identity) between genomes (observed F100) to the number expected by chance based on FastANI, allowing inference of recombination frequency within or between groups while controlling for shared ancestry ^38^.

We first examined F100 patterns across the *S. baltica* complex using all-against-all pairwise whole-genome comparisons (Fig. 5A). As expected, F100 values dropped sharply between *S. baltica* complex members and more distant species isolated from Vaxön sediments, e.g., *S. hafniensis* (SP1S2-9) and *S. oncorhynchi* (L-17, SP1S1-3), sharing less than 1% of genes [ln(F100) > 4.5] (Fig. 5B). Within the *S. baltica* complex, however, we observed unexpectedly relatively high rates of gene flow between strains representing distinct species and putative genospecies, sharing a median of 3.94% of genes (Fig. 5A, B). These values are comparable to those observed in highly cohesive species with FastANI >97%, both inside and outside the genus *Shewanella*, such as *S. algae* or *Salinibacter ruber* ^39^ (Fig. S8A, B). By group, the median proportion of shared genes in G1 was 2.73 ± 0.51% per genome, G2 had 5.95 ± 2.28%, and G3 exhibited the highest gene sharing with a median of 13.6 ± 9.84% (Fig. 5A). Between G2 and G3, the median shared gene proportion was 5.3 ± 1.32%, whereas comparisons between G1 vs. G2 and G1 vs. G3 showed a much lower median of 1.95 ± 1.49% (Fig. 5A). These results indicate that G1, despite being mostly a single species (*S. baltica*), exhibits less gene sharing than the multi-species groups G2 and G3, reflecting extensive gene flow within the *S. baltica* complex, particularly between G2 and G3. From these results we inferred that the relatively elevated inter-group exchange points to recent homologous recombination, either directly between groups or indirectly via shared genes inherited from a recent common ancestor. Indeed, Bayesian inference of the evolutionary history of *S. baltica* complex strains, based on the core proteome of the complete reference genomes (n = 32), revealed that an ancestral lineage split into two main branches, one leading to G1 and the other leading to G2 and G3 (Fig. 5C).

**Figure 5.**
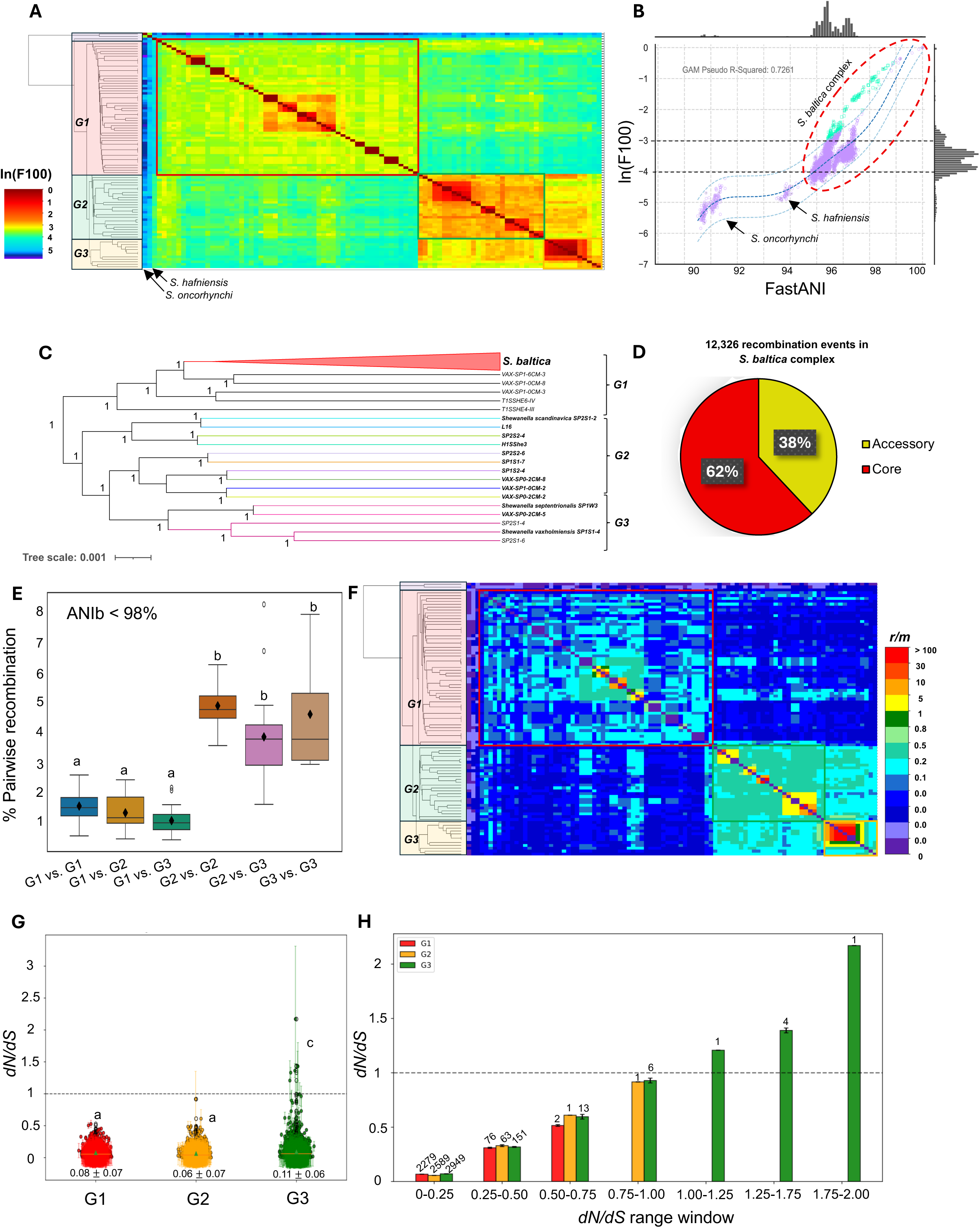
Recent recombination events within sediment strains of the *S. baltica* complex from the Stockholm archipelago. (**A**) F100 clustered heatmap showing the frequency of genes that are 100% identical between genome pairs, as a proxy of potential recombining genes. (**B**) F100 expected for their ANI range, i.e., F100 vs FastANI, building a Gam model using the sediment genomes (r^2^ = 0.72). Dashed red circle denotes the members of the *S. baltica* complex, and turquoise colors denote genomes do not follow the GAM model. (**C**) Unrooted Bayesian phylogeny tree based on core-proteome sequences of representative closed genomes of the *S. baltica* complex (n = 32). *S. baltica* subspecies were collapsed, and node labels indicate highest posterior density values. (**D**) Distribution of total recombination events (n = 12,326) across pangenome categories (core and accessory), detected from pairwise all-vs-all genome comparisons. (**E**) Percentage of recombination between pairwise genomes among the different groups within the *S. baltica* complex with ANIb < 98%. (**F**) Estimation of the ratio between the number of mutations removed through homologous recombination (*r*) and the number of mutations introduced by point mutations over the same time frame (*m*), i.e., *r/m*. In (**A**) and (**B**), F100 values were ln-transformed (ln[F100]) to improve visualization across the full range of values. In (**A**), (**B**) and (**F**) different colors represent the clades as previously described, and frames indicate the boundaries of each clade. *S. hafniensis* (SP1S1-9) and *S. oncorhynchi* (L-17 and SP1S1-3), also isolated from sediments in the same study areas, were used as outgroups for comparison. Genomes in the heatmap are ordered according to ANIb similarity. (**G**) Distribution of branch-specific *dN/dS* values across core genes in G1, G2, and G3. Boxplots show median and interquartile range, points represent individual genes, and error bars indicate standard deviation. (**H**) Mean *dN/dS* values per group across binned *dN/dS* 0.25 intervals (0-2.5), and bars represent group means within each window ± standard error, showing the number of core genes in each window for each group. In (**G**) and (**H**) the dashed line indicates neutrality (*dN/dS* = 1). For plots E and G, different letters denote significant differences among the groups (p < 0.05).

To quantify homologous recombination between these groups, we counted the genes classified as recently recombined between genome pairs. Across all genomes from our dataset, we identified 12,326 recombination events, of which 62% involved core genes and 38% accessory genes (Fig. 5D), with recombination affecting all regions of the genome (Fig. S9A). To avoid overestimating recombination due to very closely related strains, genomes with pairwise ANIb values above 98% were excluded (1.2% of the total pairwise comparisons), as these exhibited recombination levels affecting > 45% of their genome content (Fig. S9B). Among the remaining genomes, average pairwise recombination levels within G1 were ∼1.6 ± 0.5%, G1 vs. G2 ∼1.2 ± 0.6%, G1 vs. G3 ∼1.1 ± 0.5%, within G2 ∼4.9 ± 0.6%, G2 vs. G3, ∼3.8 ± 1.6%, and within G3 ∼4.6 ± 2.3% (Fig. 5E). Thus, recombination levels were significantly higher within G2, within G3, and between G2 and G3 compared with within G1 or between G1 and the other groups (p < 0.05; Fig. 5E). The length of putatively recombined segments, approximated using consecutive recombined genes, ranged from 1 to 40 kbp. For closely related genomes (ANIb > 98%), recombination segments were predominantly 1-10 kbp, whereas more distantly related genomes exhibited shorter segments, i.e., ∼1-2 kbp within G1 and ∼1-5 kbp within or between G2 and G3 (Fig. S10A–F). Overall, the extensive gene flow detected between G2 and G3 suggests recent diversification from a shared ancestry, likely driven by ‘sexual’ speciation.

We next estimated the ratio of mutations removed by homologous recombination (*r*) to mutations introduced by point mutations (*m*), i.e., the *r/m* ratio between genome pairs (Fig. 5F). Closely related genomes (ANIb > 98%) exhibited most of the *r/m* ratios ranging from 1 to 100, with a few outlier genome pairs exceeding 100 in the 99-99.5% ANIb range due to the high identity of the recombined genes identified. This pattern is consistent with high recombination rates among similar genomes and supports a role for homologous recombination in maintaining genomic cohesion among highly similar strains. By contrast, more distantly related genomes showed much lower *r/m* ratios, mostly between 0.05 and 0.2, particularly within G1 (Fig. 5F). This pattern aligns with the high internal diversity observed in G1, suggesting that point mutations predominantly drive diversification in this lineage. In G2 and G3, most *r/m* values ranged from 0.2 to 0.8 across all pairwise comparisons, showing that the relatively high recombination levels detected within and between genospecies and a diversification history can be driven by ‘sexual’ speciation.

### The *S. baltica* complex is under purifying selection

Finally, we quantified selective pressures in single-copy core genes across the different evolutionary groups (G1, G2, and G3) using branch-specific *dN/dS* (ω) estimates under the MG94 codon model (see Materials and Methods) (Fig. 5G, H). The mean ω across all groups was 0.08 ± 0.11, and approximately 99.5% of core genes exhibited ω ≪ 1 (Fig. 5G), indicating strong evolutionary constraint. Significant variation in mean ω was observed among groups (p < 0.05, Fig. 5G). The mean ω of G3 was approximately 1.5-fold higher than that of G1 and G2, with values of 0.11, 0.08, and 0.06, respectively, and G3 contained the largest number of genes with ω > 0.5, including up to 25 core genes, and a subset of six genes inferred to be under positive selection (ω > 1) (Fig. 5H), all annotated as hypothetical protein-coding genes. The overall distribution of low ω values across groups is consistent with genome-wide patterns observed in other bacterial populations ^40^, reflecting strong purifying selection acting on the core genome. These results suggest that evolutionary dynamics within the *S. baltica* complex is dominated by strong purifying selection.

## DISCUSSION

What constitutes a bacterial species and the processes underlying bacterial speciation remain as open questions that are often challenging to address empirically in natural populations. Speciation is generally understood as the emergence of genomically cohesive lineages that evolve together while gradually becoming reproductively isolated through barriers to gene flow ^7, 14^. However, such isolation is frequently incomplete, as distinct bacterial species can maintain a certain level of gene exchange through recombination ^41^. Numerous models have been proposed to describe the multifaceted modes of speciation in bacteria, which can enormously vary depending on biological and ecological factors ^20, 42–45^. The *S. baltica* complex provides a direct empirical demonstration of species concepts that view species as metapopulation lineages shaped by genetic exchange and shared ecological pressures along a continuous speciation spectrum ^18, 20, 21^. Within this complex, coexisting, yet genomically differentiated lineages incipiently diverge, likely driven by homologous recombination and ecological specialization.

In this study, we identified closely related populations comprising three distinct evolutionary and functional groups, referred to as G1, G2, and G3, respectively. Our analysis of genomic relatedness in members of the *S. baltica* complex, as inferred by all-against-all ANIb determinations, revealed a bimodal distribution. One major group of ANIb interactions, ranging from 95.6 to 97.5 ANIb% mainly involved *S. baltica* strains found in both the water column and the sediment. Remarkably, a second group of ANIb interactions fell in the range 94.5–95.6 ANIb% and was densely occupied by members of G2 and G3, encompassing numerous distinct, sediment-native genospecies according to current taxonomic delineation criteria ^34^, i.e., three species with standing in nomenclature, which we had previously described ^32, 33^, and 11 putative genospecies identified in this study. Such an ANIb distribution is remarkable, as the ANIb range preponderantly comprising interactions involving G2 and G3 members is usually scarcely populated and represents a so-called ‘gray zone’, that is, a gap separating distinct species ^4, 46^.

The analysis of G1 evolution primarily highlights *cladogenesis* at the subspecific level. This distinction is important because *cladogenesis* and *speciation* are not intrinsically synonymous. Originally introduced by B. Rensch, ‘cladogenesis’ refers broadly to phylogenetic branching and splitting ^47, 48^. In prokaryotes, a cladogenetic event does not necessarily result in the formation of two distinct gene pools that qualify as separate biological species, and it might instead produce subspecific units such as genomovars, phylogroups, or ecological subspecies. Examples include the diverse G1 ecotypes examined in this study, as well as those discussed in previous taxonomic revisions ^49, 50^, which may or may not coexist indefinitely with the parental lineage or experience ongoing gene flow. The *S. baltica* complex is notable because the genomic relatedness of its members lies near the threshold for prokaryotic species circumscription, making cladogenetic events more likely to lead to full speciation rather than mere subspecific divergence. This is illustrated in several G1 lineages that cannot be classified as coherent genospecies, and particularly in G2 and G3 members, which share a common ancestor that already diverged from the *S. baltica* lineage, as shown in our Bayesian phylogenetic reconstruction.

Speciation involves genomic differentiation and a consequent reduction in recombination between emerging lineages. The ‘porous’ nature of gene flow barriers in bacteria means that incipient species can retain substantial recombination with related lineages ^51^, as reflected in the large accessory genome of G1, low within-group similarity, and high mutation-to-recombination ratio, which drives its genomic diversity. In contrast, despite the relatively high levels of recombination between G2 and G3 (5-7%), the observed gene flow is insufficient to homogenize their genomic differentiation (which would therefore abrogate speciation) and rather point to a recent, ongoing process of sympatric speciation. This is supported by the stability over time detected for several divergent genotypes, evidenced by the repeated recovery of the same genospecies in 2021 and 2023. Evolutionary models suggest that early-stage sympatric recombination can facilitate niche adaptation by combining beneficial alleles, whereas at later stages, recombination can produce low-fitness intermediates and counteract divergence ^16, 52, 53^. Such trade-offs can be resolved by reducing the number of loci under selection or by progressively restricting recombination ^52^, being an onset of the latter likely observed between G2 and G3 members.

The speciation process experienced by G2 and G3 members is further supported by genomic signatures of distinct ecological specialization. The predominance of novel genospecies emerging from sediments suggests that a sediment-associated lifestyle promotes evolutionary divergence. Metabolic adaptation to distinct ecological niches, or to alternative strategies within shared niches, plays an important role in bacterial diversification ^54^. In surficial sediments, local variations in nutrient availability, oxygen concentration, or anaerobic electron acceptors, along with other physicochemical or biological factors, can create microenvironments favoring specific adaptations ^55^. Such fine-scale heterogeneity might remain undetected by metagenomic inference, as could be the case in this study. Regardless, rapid metabolic divergence can occur even in laboratory-recreated homogeneous environments, increasing intraspecies heterogeneity ^43, 56^. Beyond the test tube, evidence shows that genomic and gene expression variations promote ecological divergence of closely related bacteria in natural ecosystems ^57^. In the *S. baltica* complex, we identified at least three mechanisms contributing to ecological distinctiveness between groups. Firstly, genomic enrichment in resource utilization pathways differentiates G1, G2, and G3. For instance, G1 genomes were enriched in carbohydrate utilization, whereas G2 and G3 showed enrichment in denitrification and nitrogen metabolism, reflecting distinct resource use strategies in sediments. Secondly, widespread gain or loss of energy acquisition pathways, independent of isolation depth, such as sulfite reduction (highly conserved in G3), tetrathionate reduction (lost in some G1 members), or DMSO reduction (acquired and highly variable across all groups), supports ecological resource specialization in a shared ecosystem rather than selection driven by spatial isolation or environmental pressures. This could also result from stochastic gene flow followed by selection of fitness-enhancing variants. Thirdly, a substantial pool of plasmid-borne genes of unknown function, while not decisive for speciation, might provide an as-yet undefined ecological toolset for their hosts.

Collectively, these observations place the *S. baltica* complex along a continuum of lineage divergence consistent with lineage-based models of bacterial speciation (Fig. 6A-C), such as de Queiroz’s method-free concept^18^ (and related work by Hey ^19^), its subsequent adaptation by Achtman and Wagner ^20^, and the speciation-spectrum model of Shapiro and Polz ^21, 58^. In these models, populations do not form discrete, fully isolated units but behave as metapopulation lineages that maintain internal genomic cohesion while gradually reducing gene flow. Thus, species correspond to independently evolving lineages and are not necessarily phenotypically distinguishable or ecologically isolated ^20^. Divergence therefore proceeds along a continuum, from (i) interconnected intraspecific subpopulations, to (ii) closely related species (genospecies), to (iii) partially genomic differentiated species linked by genome intermediates, and ultimately to (iv) fully genomic separated species (Fig. 6A-C). Applying this model to the *S. baltica* complex, we infer that an ancestral population (G0) split into two subpopulations, G01 and G02, which subsequently followed distinct evolutionary trajectories (phase I, Fig. 6A-C). G01 gave rise to the present-day G1 population, largely dominated by *S. baltica* and characterized by low *r/m* ratios together with elevated gene gain and loss, generating extensive genomic and metabolic diversity (Fig. 6A-C). In contrast, G02 diversified into the G2 and G3 lineages, which represent incipient, independently evolving species (genospecies) that remain internally cohesive yet increasingly differentiated from one another (phase II, Fig. 6A-C). Their persistence over multiple years (meaning successful ecological adaptation) and ongoing homologous recombination, indicates a metapopulation structure in which divergence is actively occurring. As recombination declines between G2 and G3, genomic isolation increases and divergence continues, generating genomic intermediates (phase III, 6A-C). Notably, *S. septentrionalis*, despite belonging to the *S. baltica* complex as a G3 representative, appears to be more closely related to *S. hafniensis* than to *S. baltica* (Fig. 5B), consistent with a possible placement of S. hafniensis in phase III of the speciation continuum (phase III, 6A-C). Finally, continued divergence and a further decline in recombination could give rise to fully genomically isolated species (phase IV, Fig. 6A–C), with *S. oncorhynchi* as a putative example (Fig. 5B).

**Figure 6.**
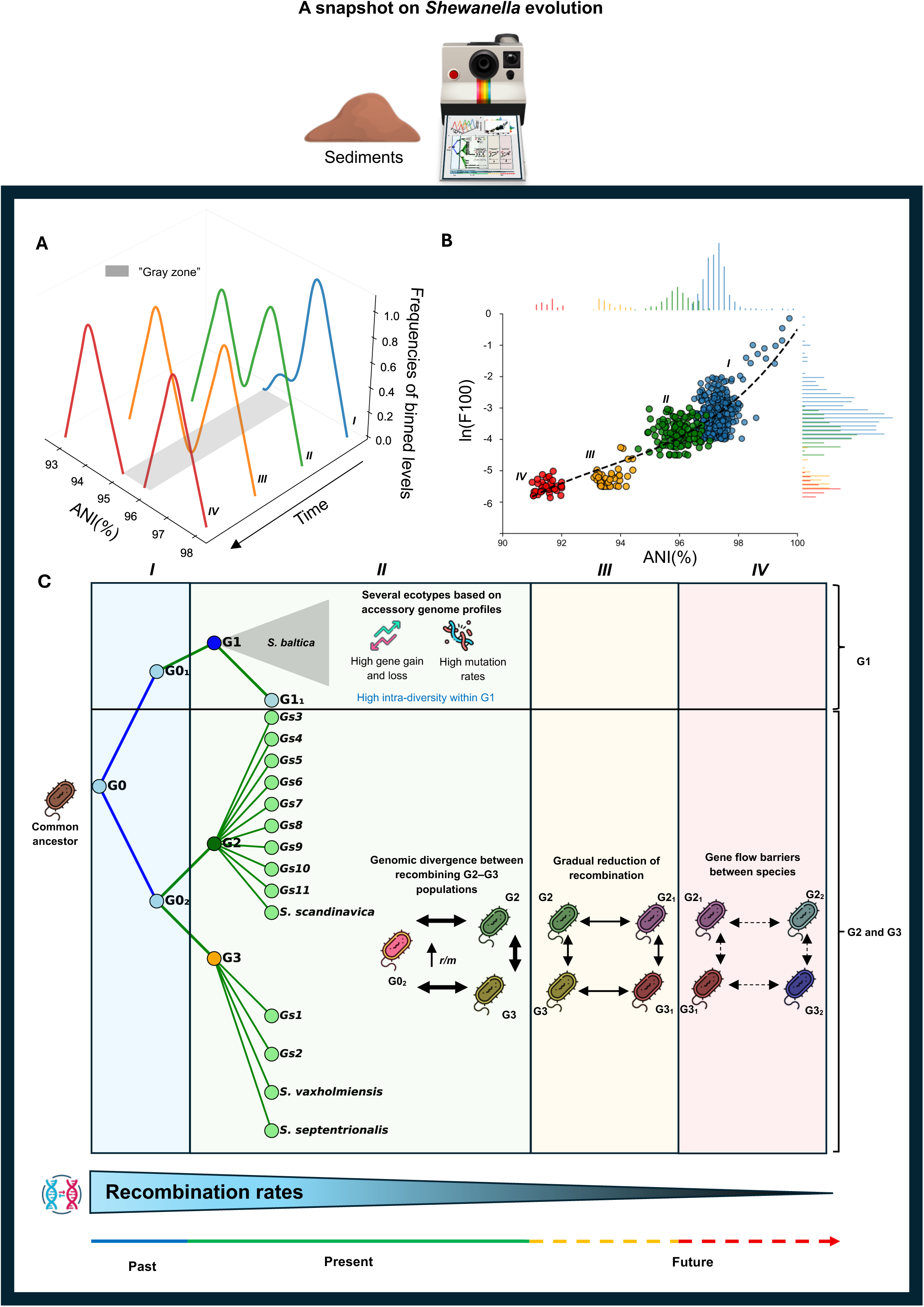
Speciation model within the *S. baltica* complex. (A) Conceptual model adapted for the *S. baltica* complex from Achtman and Wagner ^20^, (B) recombination dynamics within the complex, and (C) evolutionary trajectories of the *S. baltica* complex, showing how lineages progressively diverge through four phases from a common ancestor. In phase I (blue), high recombination maintains genomic cohesion within the ancestral lineage (G0), while subpopulations gradually emerge and begin evolving independently (G01 and G02). In phase II (green), recombination rates decline but remain substantial. During this stage, G01 gave rise to the G1 population (predominantly *S. baltica*), characterized by low *r/m* ratios and high gene turnover, whereas G02 differentiated into the G2 and G3 lineages, representing incipient, independently evolving species (genospecies) whose divergence is likely driven by homologous recombination. Together, ongoing recombination and the recurrent detection of some these lineages support their gradual genomic and ecological differentiation. In phase III (orange), recombination decreases further, resulting in increasing genomic isolation between lineages. In phase IV (red), lineages become fully differentiated species with minimal recombination and effective genomic isolation.

In conclusion, the extent of diversity uncovered in the sediment populations analyzed here has not previously been observed among *S. baltica* strains isolated from the stratified Baltic Sea water column ^28, 29^ or from other environments, pointing to a sediment-associated lifestyle as a potential catalyst for the emergence of novel *Shewanella* genotypes through recombination-driven (‘sexual’) speciation and metabolic specialization. Support for this view is provided by the documented coexistence of multiple recently described *Shewanella* species in mangrove surficial sediments, whose close genomic relatedness to *S. loihica* mirrors the diversification patterns observed within the *S. baltica* complex ^59^. Together, these findings capture a snapshot of early-stage sympatric speciation, revealing the coexistence of incipient yet genomically differentiated lineages within a shared habitat lacking obvious spatial barriers, where the intrinsic lifestyle in this environment appears to actively promote diversification. Future work should determine how sediment microhabitats function as crucibles for speciation, elucidate the mechanisms driving divergence under continued gene flow, and identify the ecological determinants that sustain the coexistence and persistence of emerging genotypes.

## MATERIALS AND METHODS

### Sample collection, environmental DNA extraction, and bacterial isolation

Between 2021 and 2023, surficial sediment cores were aseptically collected at Vaxön (Stockholm archipelago, 59°24′01.0″ N 18°20′20.2″ E) using sterile 30 ml syringes (VWR Cat. No. 613-2035). Samples were transported to the laboratory within 2 h, and aliquots for DNA extraction and bacterial isolation were taken at 0, 2, 4, and 6 cm depth. DNA (ca. 0.5-0.8 mg) was extracted using the DNeasy PowerSoil Pro kit (Qiagen, Hilden, Germany), after mechanical disruption with FastPrep-24™ 5G (MP Biomedicals; 4 × 40 s at 5.5 m/s) and stored at −80 °C until sequencing. For bacterial isolation, sediment aliquots were suspended in phosphate-buffered saline (0.01 M, pH 7.4), vortexed, serially diluted, and plated (100 µl) on Lyngby’s Iron Agar (LIA) with 0.04% w/v L-cysteine. Plates were incubated at 28 °C for 24 h. *Shewanella*-like colonies, characterized by black-centered with pale orangish margins, were re-streaked three times on LIA and identified to the genus level by MALDI-TOF (MALDI Biotyper Sirius, Bruker). Additional sediment and water samples were collected from Hagaparken (59.3546 N, 18.0429 E), Lidingö (59.3667 N, 18.1757 E), Nynäshamn (58.8995 N, 17.9508 E), and Tyresö (59.2324 N, 18.3110 E) between 2021 and 2022 (Fig. S1), and processed as previously described ^32, 33^. In total, 112 *Shewanella* spp. strains were isolated and analyzed, including previously reported strains ^32, 33^.

### Whole-genome and metagenome DNA sequencing

Genomic DNA was extracted from overnight LB (Miller) cultures at 28 °C using the DNeasy Blood & Tissue kit (Qiagen) and quantified with the Quant-iT High Sensitivity dsDNA assay (ThermoFisher, Waltham, MA, USA). Equimolarly pooled libraries were circularized using the MGI Easy Circularization kit, and 2×100 bp paired-end sequencing was performed on a DNBSEQ-G400 instrument (MGI, Shenzhen, China). Genome assemblies were generated with SPAdes v3.15.5 ^60^ via Bactopia v1.7.x^61^ or BACTpipe (https://github.com/ctmrbio/BACTpipe). For 32 representative strains, including type strains and at least one novel genospecies, Oxford Nanopore sequencing was performed to obtain complete genome sequences. High-molecular-weight DNA was extracted using the Quick-DNA HMW MagBead Kit (Zymo Research, Irvine, CA, USA), and libraries were prepared with the Rapid Barcoding Kit (SQK-RBK114.24). Sequencing was carried out on a FLO-MIN114 (R10.4.1) flow cell using a MinION Mk1B device (Oxford Nanopore Technologies), using MinKNOW v25.05.14 with high-accuracy basecalling modelling. Long reads were assembled *de novo* using Flye v2.9.5-b1801 ^62^ and Raven v1.8.3 ^63^. Hybrid assemblies combining Oxford Nanopore long reads and DNBSEQ-G400 reads were generated using Unicycler v0.5.1 to optimize plasmid recovery ^64, 65^. Circularized replicons produced by Flye were retained when possible, and genomes were polished using Medaka v1.11.3 (https://github.com/nanoporetech/medaka), Polypolish v0.6.0 ^66^, and POLCA within MaSuRCA v4.1.0 ^67, 68^, and reordered using UGENE v48.1 ^69^ based on the Unicycler start genes database ^65, 69^. Assembly quality metrics were evaluated using QUAST v5.0.2 ^70^. Genome completeness and contamination were evaluated using CheckM^71^ and ConFindr^72^. The genomes were of high quality, exhibiting an average of 6 ± 4 single nucleotide variants (SNVs) in core genes, with mean completeness of 98.76 ± 0.86% and contamination of 0.74 ± 0.63%, indicating a low likelihood of chimeric assemblies (Table S2).

For metagenomic analysis, 2×150 bp paired-end sequencing was performed on a DNBSEQ-T7 (MGI). Quality-trimmed reads were classified with Kraken2 v2.1.0 ^73^ using the Genome Taxonomy Database (GTDB, downloaded 8 April 2022) ^74^. Contigs were assembled with MEGAHIT v1.2.9 ^75^ and analyzed with the SqueezeMeta v1.7.2 pipeline ^76^. Open reading frames (ORFs) were predicted with Prodigal v2.6.3 ^77^, and protein similarity searches were performed against the GenBank non-redundant (nr) protein and Kyoto Encyclopedia of Genes and Genome (KEGG) databases using DIAMOND (v2.1.10) ^78^. Finally, reads were mapped back to contigs with Bowtie2 v2.5.4 ^79^ to assess functional coverage and abundance.

### Genomic relatedness analyses

Average nucleotide identity (ANI) analyses were performed using JSpeciesWS (for BLAST-based ANI (ANIb)) ^80^ and FastANI v1.34 (‘Many to Many’ mode, default settings; https://github.com/ParBLiSS/FastANI). Pairwise ANIb values were hierarchically clustered to define genome groups. Publicly available genomes of *Shewanella algae* (n = 219), *S. xiamenensis* (n = 98), and *S. oncorhynchi* (n = 32, including three strains from this study) were downloaded from NCBI GenBank on 31 May 2025 and analyzed with FastANI. Digital DNA–DNA hybridization (dDDH) values were calculated using the Genome-to-Genome Distance Calculator (GGDC) ^81^, comparing each genome against the maximum allowed number (n = 75), and repeated for all genomes. Only results from the recommended Formula 2 were reported ^82^.

### Pangenome analyses, accessory genome clustering and gene gain-loss dynamics

Pangenome analyses within the *S. baltica* complex (and the additional datasets previously mentioned) were performed on genomes annotated with Prokka v1.14.6 ^83^, using PPanGGOLiN v2.2 ^84^. Default parameters were applied with ≥90% identity and coverage, as higher thresholds (>95%) over-split conserved genes and lower thresholds (50–80%) merged paralogs. Genes present in ≥95% of genomes were classified as core, considering that the vast majority of the genomes sequences are draft quality. Core gene and protein family alignments were generated using fast Fourier transform (MAFFT) ^85^, and phylogenomic trees were reconstructed with FastTree v2.1.10 ^86^ using GTR+Γ4 for DNA alignments and JTT+CAT for protein alignments. Trees were bootstrap-supported with 1,000 replicates, midpoint-rooted, and visualized in iTOL v6 6 ^87^. The *gene_presence_absence.Rtab* matrix from PPanGGOLiN was used to construct a hierarchical clustering dendrogram based on gene presence–absence. This matrix was also analyzed with Mandrake v1.2.2 to generate a two-dimensional embedding ^88^, enabling identification of accessory genome clusters (AGCs). Gene gain and loss events were modeled using Panstripe v0.3.1, using the included Gaussian distribution, which provides more robust model fitting while yielding results comparable to the compound poisson (Tweedie) model ^36^.

### Estimation of genome-wide differentiation

Genome assemblies were aligned with Split K-mer Analysis V2 using *S. baltica* CECT 323^T^ as the reference, and single nucleotide polymorphisms (SNPs) were extracted with SNP-sites v2.5.1 ^89^ and stored in VCF format. SNPs were grouped into overlapping 10 kb sliding windows (2.5 kb step), and for each window, pairwise differentiation was calculated as the proportion of discordant SNPs (p-distance). Genome-wide differentiation between each pair of genomes was obtained as the average p-distance across all windows, providing a genome-wide complement to classical population-genetic metrics ranging from 0 (identical genomes) to 1 (completely different) ^90^. Pairwise distances were concatenated and clustered as for ANIb.

### Bayesian phylogenetic analysis

Bayesian phylogenetic analysis was performed with BEAST X v10.5.0 ^91^ using the core proteome alignment of 32 complete genomes (3,396 proteins; 1,157,272 amino acids) representing subspecific groups of *S. baltica* and related genospecies. A Blosum62 substitution model was applied, with a strict molecular clock to assume uniform substitution rates, and no time calibration was used, as the aim was to assess relative relationships rather than absolute divergence times ^92, 93^. The starting tree was generated under a constant-size coalescent, and a Yule speciation model constrained tree topology and branch lengths. Posterior distributions were estimated via Markov Chain Monte Carlo (MCMC) for 10,000,000 generations, sampling every 1,000 steps. Convergence was assessed with Tracer v1.7.2 ^91^, trees were summarized with TreeAnnotator v10.5.0 ^91^, and visualized using iTOL v6 ^87^.

### Metabolic and horizontally acquired gene profiling for ecological differentiation

In KBase ^94^, genome assemblies were re-annotated using RAST SEED roles ^95^ by calculating the proportion of protein-coding genes assigned to each function. Genes within regions of genome plasticity (RGPs), defined by PanRGP as clusters of consecutive shell or cloud genes ^96^, were used as a proxy for horizontal gene transfer (HGT). These genes were annotated with COGclassifier v1.0.5 based on COG categories (https://github.com/moshi4/COGclassifier/tree/main), and taxonomic assignments were obtained using EggNOG v5.0 (Evolutionary Genealogy of Genes: Non-supervised Orthologous Groups) ^97^. Functional enrichment was assessed using Kruskal-Wallis tests for SEED-role and HGT-like genes COG categories, and fold changes were calculated as the median value per genome relative to all others. Taxonomically annotated metagenomes, RAST-annotated genomes, and COG-annotated HGT genes were compiled into feature matrices and standardized to Z-scores prior to multivariate analyses, including principal component analysis (PCA). Group differences were assessed using permutation-based multivariate analysis of variance (PERMANOVA), and enrichment analyses used Benjamini–Hochberg false discovery rate (FDR) correction, with significance defined as FDR-adjusted p < 0.05 and fold-change thresholds of > 1.2 or < 0.8. Finally, KEGG pathway module completeness was evaluated from KEGG orthologues (KOs) presence-absence using the kegg-pathways-completeness-too (https://github.com/EBI-Metagenomics/kegg-pathways-completeness-tool/tree/master).

To investigate the respiratory metabolism of N- and S-oxides, we first compiled annotated sequences of respiratory reductases from previously described species within the complex, including *S. baltica* OS195 (GCA_000018765.1), *S. baltica* CECT 323^T^ (NZ_UGYM01000002.1), and *S. scandinavica* SP2S1-2^T^ (JAUOES000000000.1). These genomes contained reductases for nitrate (NapA; ABX48036, ABN61416, MDT3282522, MDT3279942, MEM6250911, MEM6249282), nitrite (NrfA; ABX50994, MDT3279623, MEM6250231.1), trimethylamine *N*-oxide (TMAO, TorA; ABX50514, MDT3281430.1, MEM6250556.1), dimethyl sulfoxide (DMSO, DmsA; ABX49397.1, MDT3282918.1), tetrathionate (TtrA; WP_006080095, MDT3279598.1, WP_311905661), thiosulfate (PhsA; MDT3282209.1, WP_311905956), and sulfite (SirA; ABX51162, MDT3279461, MEM6249422). After identifying homologs in our dataset, we constructed a custom database of these respiratory reductases designed to capture the diversity of evolutionary lineages within the complex. For the final analysis, only the curated reference sequences from this database were used for BLASTp (BLAST+ v2.1.16) ^98^ searches with an e-value cutoff of 10^−30^, and hits with ≥90% sequence identity were retained for downstream analyses.

### Detection of recent recombination events

Recent recombination was assessed using the F100 prokaryotic recombination pipeline ^38^ (https://github.com/rotheconrad/F100_Prok_Recombination). Reciprocal best matches (RBMs) between genome pairs were identified by BLAST ^99^, and the F100 metric was calculated as the proportion of RBMs that were 100% identical, serving as an indicator of recent gene exchange. Recombinant genes were classified as core (≥90% of genomes), accessory (<90% and present in multiple genomes), or genome-specific, while highly conserved genes (as the 10% of core genes with the lowest divergence) were excluded to avoid conflating conservation with recombination. Gene clustering was performed with MMSeqs2 using the same parameters as for core gene definition (--min-seq-id 0.90, -c 0.90, --cov-mode 1, --cluster-mode 2, --cluster-reassign), and pairwise genome recombination values were extracted from the pipeline output (script *03f_Recombinant_pair_analysis.py*, default settings).

### Evolutionary genomic analyses

The relative contribution of homologous recombination to point mutation was quantified using the *r/m* ratio, defined as the number of nucleotide differences removed by recombination (*r*) relative to those introduced by mutation (*m*). Pairwise *r/m* estimates were obtained using the empirical approach implemented in the F100 prokaryotic recombination pipeline ^38^, which assumes that recombined genes initially carried the average sequence divergence of the compared genomes.

Selection acting on core genes of the *S. baltica* complex was assessed using codon alignments of single-copy core genes for each respective phylogenetic group described in this study. Protein alignments were back-translated to codon alignments using PAL2NAL v14 ^100^ to preserve reading frame information. Gene-specific phylogenies were reconstructed with FastTree v2.1.11, and branch-specific ratios of nonsynonymous to synonymous substitution rates (*dN/dS,* ω) were estimated using the FitMG94 model, which implements the Muse–Gaut (MG94) codon substitution model of sequence evolution, in HyPhy v2.5.64 ^101^. The model was implemented to permit independent variation of *dN* and *dS* across branches, thereby accommodating lineage-specific heterogeneity in selective pressures. Confidence intervals for ω were estimated using profile likelihood, and deviations from neutrality (ω = 1) were evaluated using likelihood ratio tests comparing the unconstrained model to a nested model in which ω was fixed at 1. To ensure robust downstream inference, HyPhy outputs were filtered to retain genes with codon alignments of at least 60 codons, and estimates were excluded when *dS* were below 0.005 to minimize instability and avoid artificially inflated ω values associated with extremely small *dS* values. For core genes showing signatures of positive selection, we additionally screened for homologous recombination, and removed if it was the case, using the GARD method ^102^ implemented in HyPhy, since recombination can produce false signals of adaptive evolution by affecting *dN/dS* estimates.

### General computational methods, stats, and data availability

Data processing, plotting, and statistical analyses were performed using custom Python (v3.9.19) and Bash (GNU Bash v3.2.57) scripts, together with RStudio v2025.05.1. All analysis scripts used in this study available via the Martín-Rodríguez Lab GitHub repository (https://github.com/Martin-Rodriguez-lab/Life-in-sediments-fosters-sexual-speciation-in-Shewanella-baltica), and multiple sequence alignments and datasets used and/or analyzed to reproduce the analysis can be found in our Figshare for this project (10.6084/m9.figshare.30902399). Genome assemblies are available in NCBI under BioProject PRJNA526057 and metagenomic raw reads under SRR36813638-SRR36813649, with individual accession numbers listed in Table S2 within File S1.

## ACKNOWLEDGEMENTS

Research at the Martín-Rodríguez laboratory was supported by grants from the Spanish Ministry of Science, Innovation and Universities (Agencia Estatal de Investigación, Ref. PID2023-150655NA-I00), the University of Las Palmas de Gran Canaria (Ref. CEI2022-04), the Anna & Gunnar Vidfelts Foundation for Biological Research (Ref. 2024-169-Vidfelts Fond), and the Swedish Research Council for Sustainable Development (FORMAS, Ref. AC-2023/0032). Bioinformatic computations were partially performed on resources provided by the National Academic Infrastructure for Supercomputing in Sweden (NAISS) at Dardel.

## AUTHOR CONTRIBUTIONS

**VF-J:** Data curation, formal analysis, investigation, methodology, software, validation, visualization, writing – original draft, writing – review and editing. **FS-S:** Formal analysis, methodology, software, writing – review and editing. **GS:** Formal analysis, methodology, software, writing – review and editing. **AJM-R:** Conceptualization, data curation, funding acquisition, investigation, methodology, project administration, resources, supervision, validation, writing – review and editing.

## SUPPLEMENTARY IINFORMATION

**Figure S1. Geographic locations of sampling sites in the Stockholm region.** The number of isolates collected at each site and used in this study (n = 113), and an overview of the experimental design used in this study. Across the paper the following abbreviations are used refer to the sampling sites; Växon 2023, VAX-SP (59.3999 N 18.3374 E); Växon 2021, SP (59.401044 N 18.325925 E); Hagaparken, H (59.355833 N 18.043611 E); Lidingö, L (59.367700N 18.156943 E); Tyresö, T; Nynäshamn, N (58.899247 N 17.950619 E). The size of the circle denotes the numbers of isolates from each sampling site.

**Figure S2. Taxonomic and functional profiles of metagenomic samples collected from different sediment depths. A**) Top 15 most abundant phyla with the Shannon diversity, and **B**) beta-diversity analysis using principal component analysis (PCA) to assess genus-level dissimilarity between the different depths in Växon sediments. Analyses were conducted on classified reads using Kraken2 and Bracken2 against the GTDB database downloaded on April 8, 2022. (E) PCA and (**D**) heatmap of gene abundance per million (GPM) annotated contigs, derived from KEGG annotations, and **E**) the top 100 most abundant KEGG annotations-based GPM, categorized by functional class found across all the collected depths, with average values log_10_-transformed. The assembled contigs were KEGG annotated using the SqueezeMeta pipeline. **F**) Relative abundance of *Shewanella* in the metagenomic bacterial reads.

**Figure S3. Genomic diversity of the *S. baltica* complex based on ANI and shared genome fraction.** (**A**) Frequency distributions of pairwise ANIb comparisons (0.1% windows) for *S. baltica* complex genomes (n = 109), including *S. hafniensis (*SP1S1-9 and 3 additional from NCBI, Table S3) *and S. baltica* CECT 323^T^ *(top red histogram), in combination with a* pairwise comparisons (blue dotplot), for showing the share gene content (**B**) Kernel density distribution of FastANI values, with distinct peaks corresponding to *S. baltica* (blue; genomes annotated as *S. baltica*, n = 70, in this study plus publicly available genomes annotated as *S. baltica,* n = 66), *S. xiamenensis* (green), *S. oncorhynchi* (yellow), and *S. algae* (red). All genomes were downloaded from the NCBI GenBank on 31 May 2025 (Table S3).

**Figure S4. Core genome-based and core proteome-based phylogenomic reconstructions of the *S. baltica* complex.** (A) Combined core genome and proteome tree constructed using a 90% identity and 90% coverage threshold based 3,308 core gene and protein families from the 109 genomes analyzed. The type strain *S. baltica* CECT 323^T^ was included as a reference. Blue labels indicate genomes isolated from the water column. Within G2 and G3, additional colors indicate distinct species previously described, namely *S. scandinavica*, *S. vaxholmiensis*, and *S. septentrionalis*, as well as putative genospecies identified in this study. The species delineation is explained in the main text. “Gs.” followed by number in the plot means putative genospecies identified in the *S. baltica* complex, and numbers on the branches denote bootstrap support values after 1,000 replications. (**B**) Accumulation curves of accessory genome content of the *S. baltica* complex compared with the publicly available genomes of *S. xiamenensis* (green), *S. oncorhynchi* (yellow), and *S. algae* (red). All genomes were downloaded from the NCBI GenBank on 31 May 2025.

**Figure S5. Regions of genome plasticity (RGPs) in the *S. baltica* complex.** (A) Boxplot illustrating the distribution of RGP counts per genome within each group. (B) Taxonomical classification of genes detected within RGPs, assigned using EggNOG, across the different groups. No significant changes were found across the groups for (**A**) and (**B**) (p > 0.05).

Figure S6. Completeness of KEGG metabolic pathways. KEGG pathways were annotated for the defined groups to assess the completeness of metabolic routes across the study strains within the *S. baltica complex*. Carbohydrate-related pathways, highlighted with arrows, were more complete in Group 1 (G1).

**Figure S7. Plasmid characterization within the *S. baltica species* complex.** (**A**) Pairwise average FastANI comparison of all plasmids identified within the *S. baltica* complex among the 32 closed genomes. Numbers after the name denote the plasmid number in that specific genome. (**B**) Cluster of orthologous groups (COG) annotation of the 21 detected plasmids. Detailed gene annotations are provided in Table S7.

**Figure S8. F100 vs FastANI** of (**A**) *S. algae* (n = 219) and (**B**) *Salinibacter ruber* (n = 102) from a saltern in Mallorca. The dashed black lines indicate the ln(F100) interaction boundaries between ln(F100) = −4 and ln(F100) = −3. For (**A**) and (**B**), a Gam model using the public available genomes from NCBI (downloaded 31 May 2025) was built (r^2^ = 0.72), and turquoise colors denote genomes that do not follow the model.

**Figure S9. Reciprocal best match (RBM) genes identified for several *S. baltica complex***. (**A**) Pairwise RBM genes were identified across several genomes within the *S. baltica* complex, including VAX-SP0-4CM-5 (G1), VAX-SP1-2CM-2 (G1), *S. scandinavica* SP2S1-2 (G2), SP1S1-7 (G2), *S. vaxholmiensis* SP1S1-4 (G3), and S2S1-4 (G3). Each rectangular marker in the figure represents an individual gene, color-coded to distinguish highly conserved/universal genes (in red), core genes (in blue), and accessory genes (green). The y-axis indicates nucleotide sequence identity of RBM genes shared between the seven query genomes (each shown as a separate row) and the reference genome (x-axis), where genes are ordered by their position in the reference genome. Green arrows indicate genomic islands unique to the reference genome, characterized by the absence of corresponding genes in other genomes in the same region. In contrast, red arrows point to highly conserved regions consistently shared within the pairwise. (**B**) Recombination comparison between genomes from different groups (G1, G2, and G3) of the *S. baltica* complex belonging to the same species, and ANIb > 98 %. Asterisks indicate functional categories significantly enriched among recombinant genes, as determined by one-sided Chi-square tests (*p* < 0.05) with Benjamini-Hochberg correction, suggesting these functions might be preferentially transferred via homologous recombination.

**Figure S10. Length distribution of recombinant fragments among genomes**. These graphs illustrate the frequency and distribution of consecutive recombinant genes, serving as a proxy for recombinant fragment lengths, as well as the distribution of consecutive non-recombinant genes. Analyses were performed across representative genomes within the *S. baltica* complex, including VAX-SP1-2CM-2 (G1), L-3 (G1), *S. scandinavica* SP2S1-2 (G2), SP1S1-7 (G2), *S. vaxholmiensis* SP1S1-4 (G3), and S2S1-4 (G3). Similar results were obtained with other pairs of genomes (not shown) among the different members of the groups described.

**Table S1. Metagenomic summary and taxonomic composition of VAX soil samples across treatments (SP0-SP4) and depths (0–6 cm).** For each sample, total raw reads, percentages of classified, unclassified, and microbial reads are reported. The relative abundance (%) of dominant bacterial genera is shown, with corresponding taxonomic lineage.

**Table S2. Metadata and genome assembly statistics for the isolates used in this study.** Genomes were annotated using Prokka, and genome completeness and contamination were assessed with CheckM.

**Table S3. Public genome sequenes from NCBI GenBank included in this study**. Assembly accession numbers for representative strains of *S. baltica*, *S. algae*, S*. oncorhynchi*, and *S. xiamenensis* used for comparative genomic analyses are listed.

**Table S4. SEED-roles. Differentially abundant SEED functional roles identified between groups.** For each role, the accessory group, comparison group, adjusted p-value, and fold change are reported.

**Table S5. Differentially abundant COG-annotated genes between groups.** For each gene cluster, the COG identifier, functional category (COG letter), group assignment, accessory gene cluster number, fold change, and predicted gene product are reported.

**Table S6. Identification of genes encoding reductases for alternative electron acceptors across study genomes.** Query sequences were matched to group-specific reference genes using sequence similarity searches, and alignment statistics (% identity, alignment length, mismatches), together with genome, group assignment, and sampling source, are reported.

**Table S7. Plasmid annotations for study genomes.** For each plasmid, the number and size (Kb), associated clade, coding sequences (CDSs), genomic location, and predicted gene product are listed.

