## Supplementary figures and images for "Life in sediments fosters ‘sexual’ speciation in *Shewanella baltica*"

### Figure S1

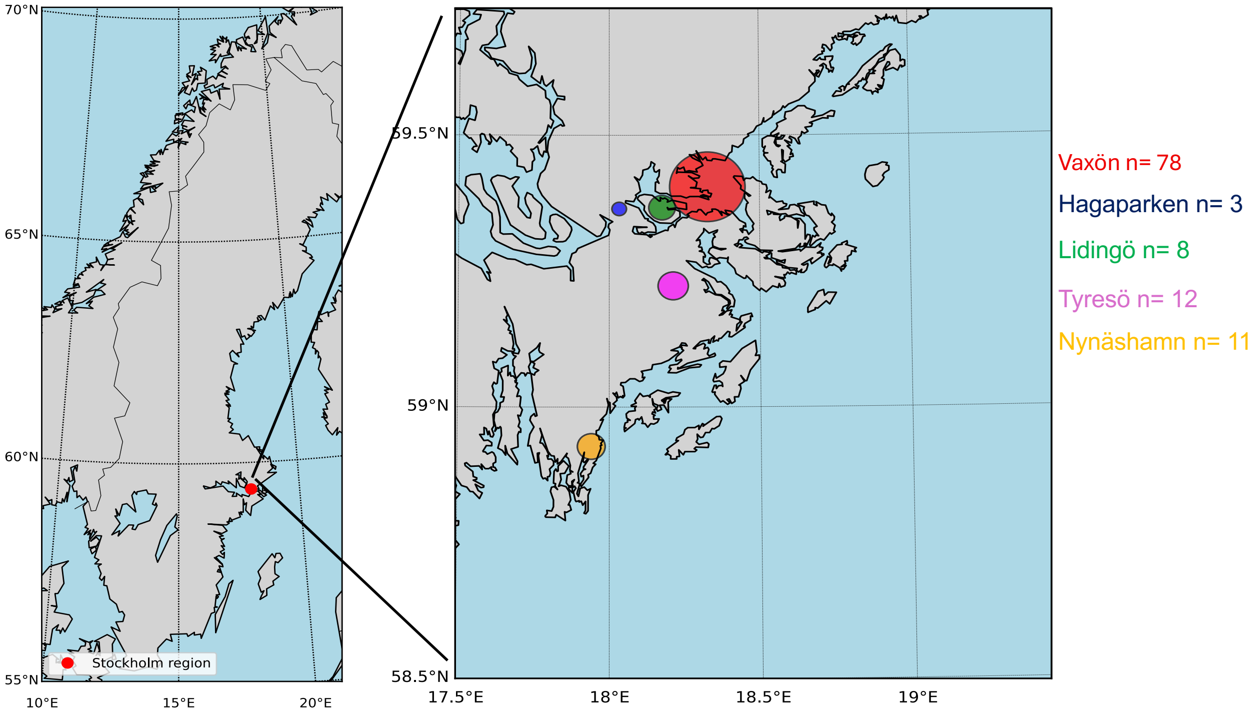

### Figure S2

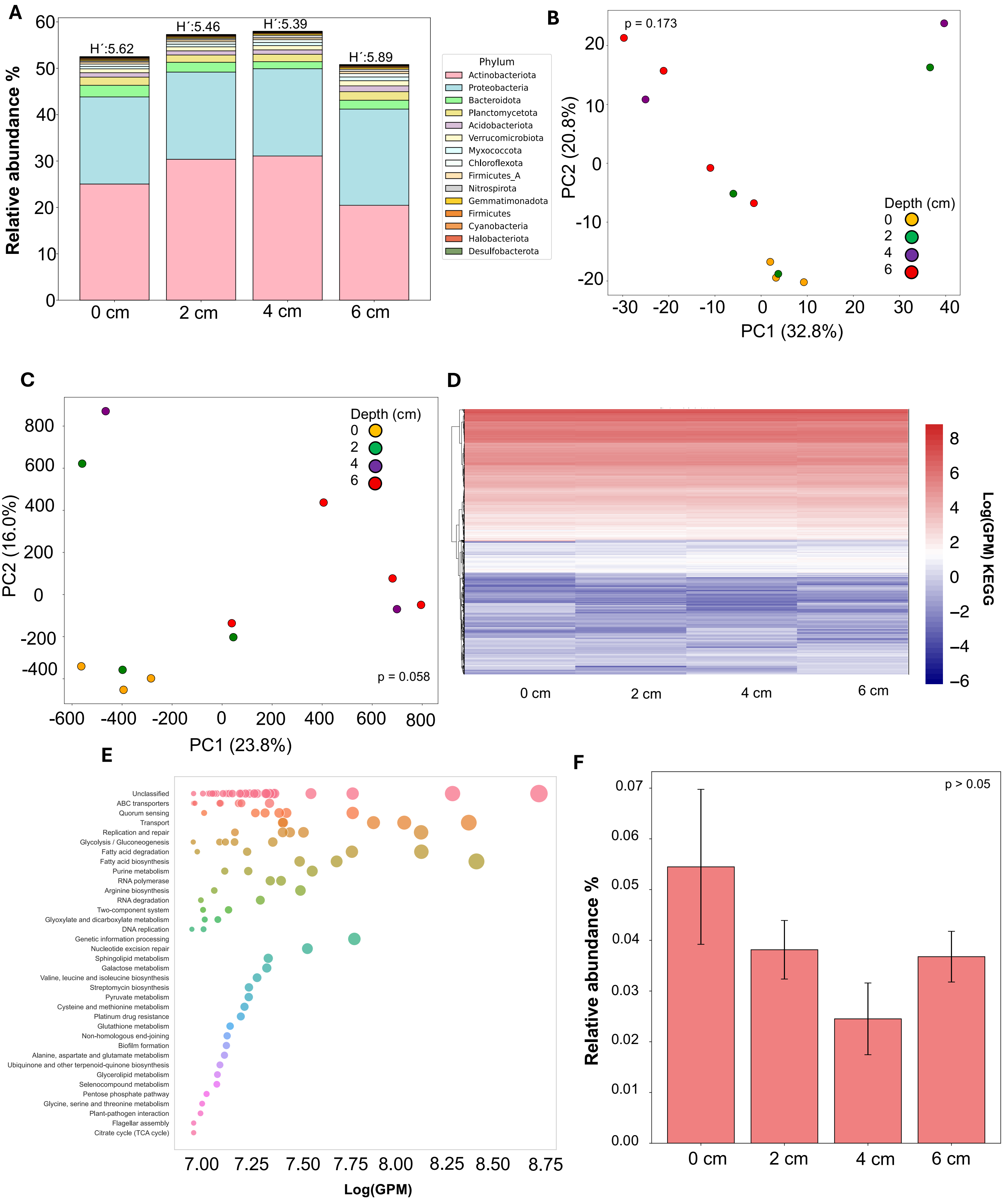

### Figure S3

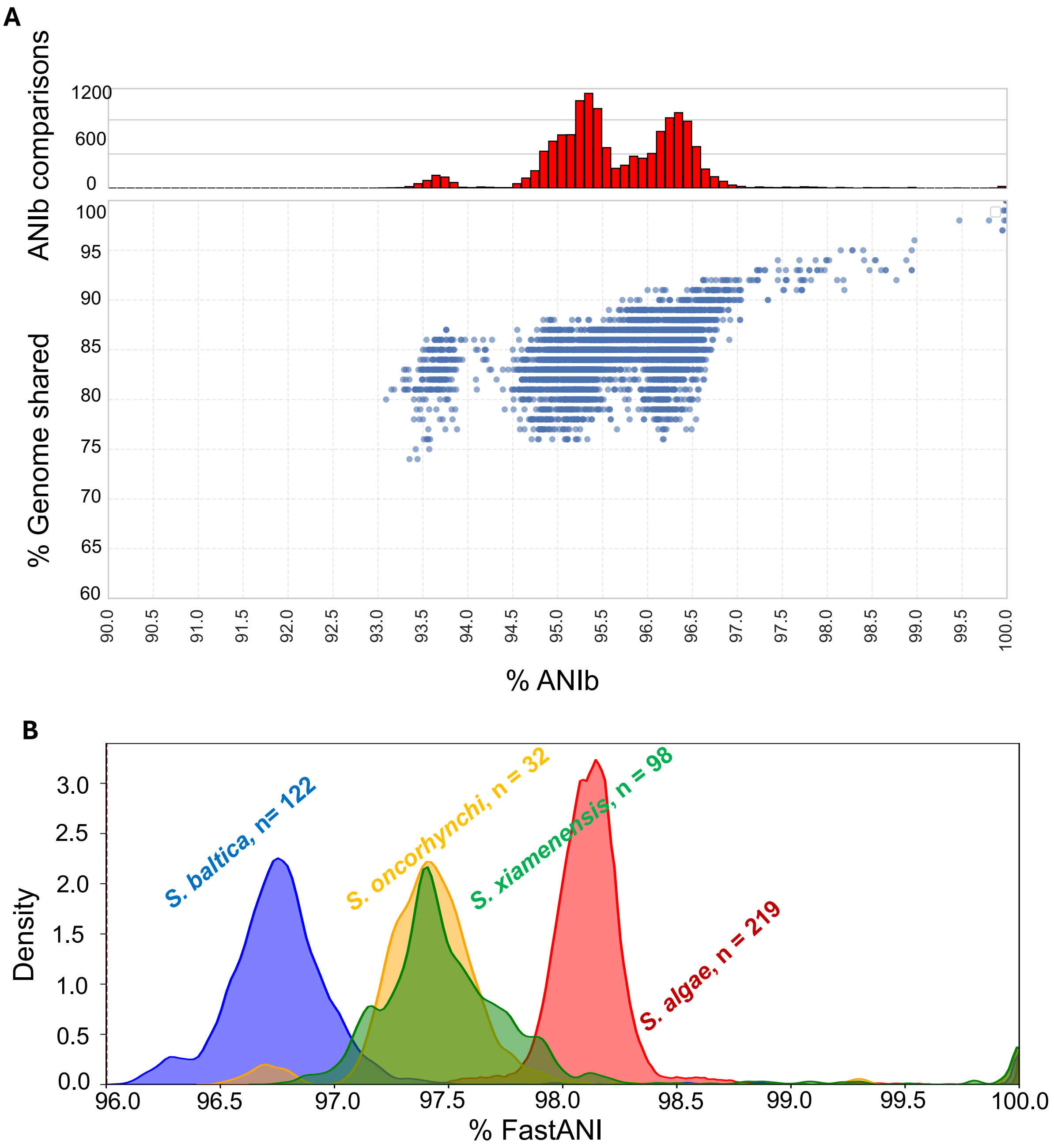

### Figure S4

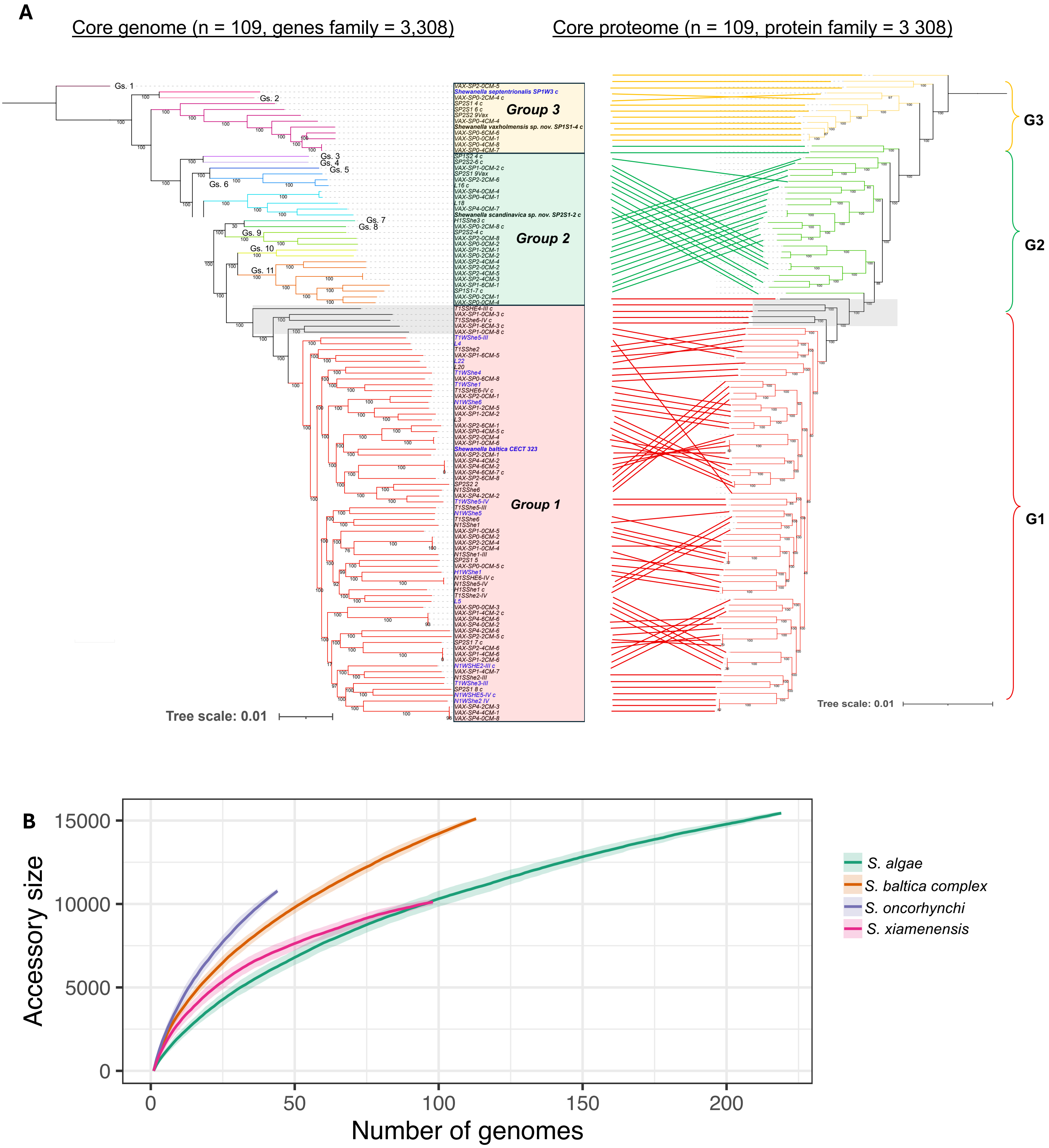

### Figure S5

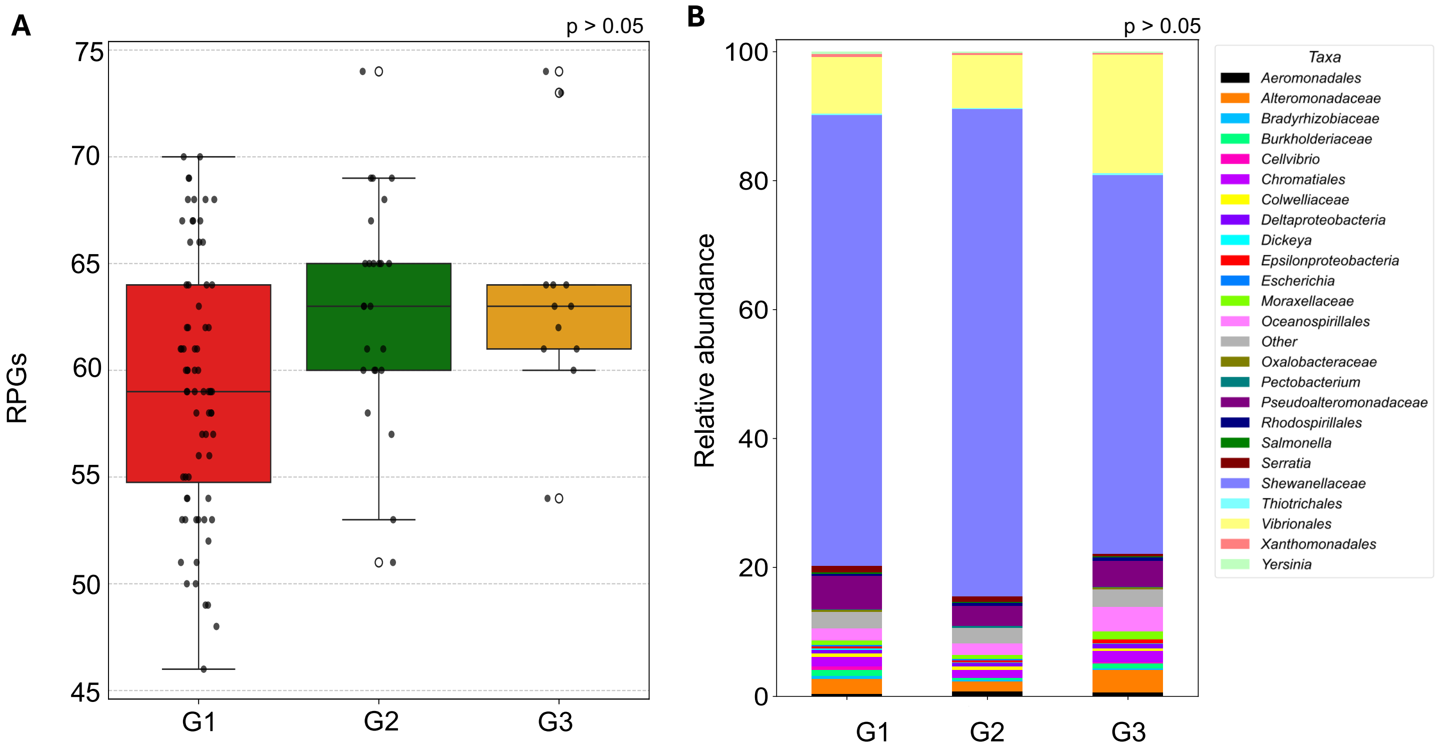

### Figure S6

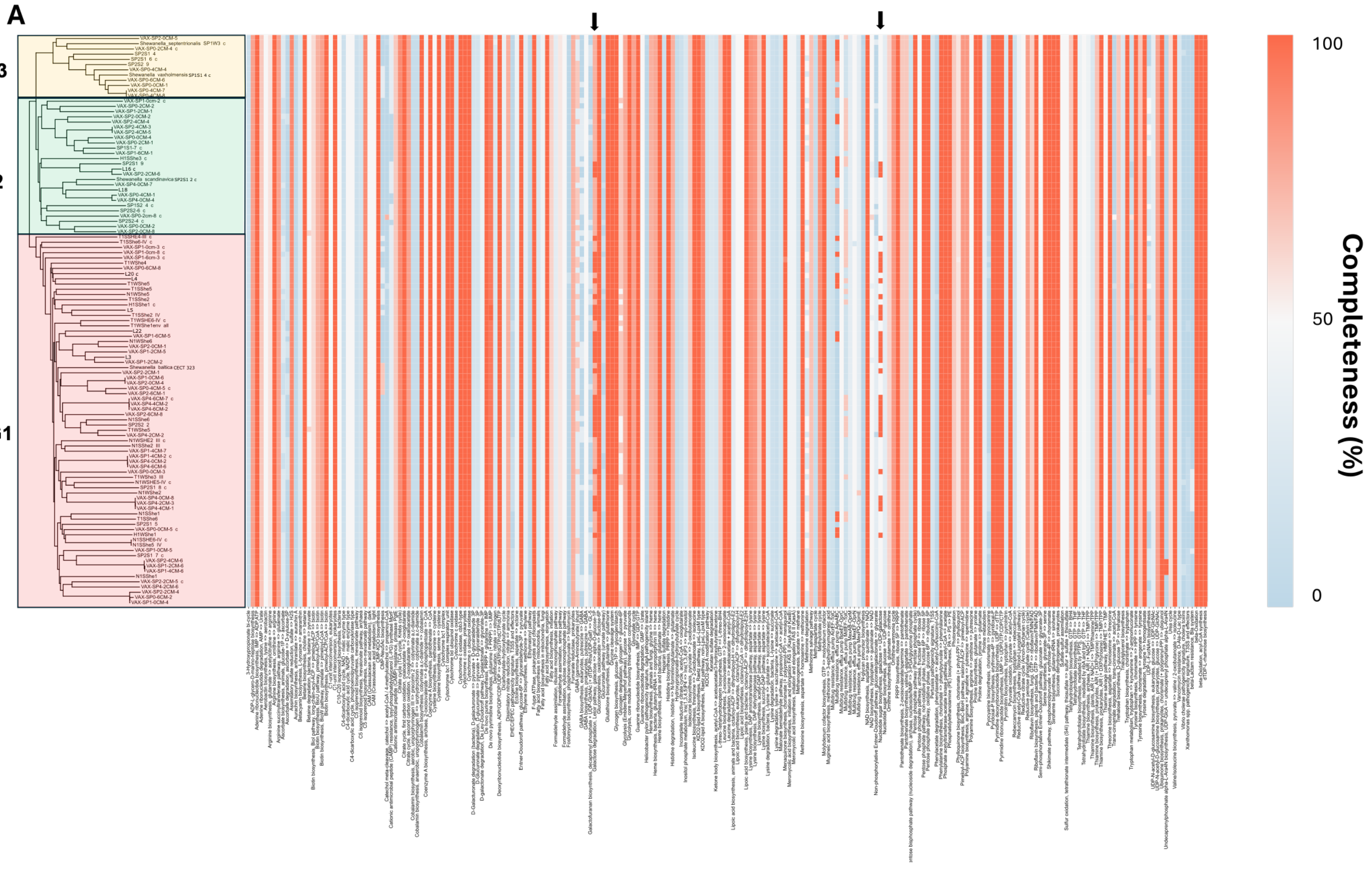


**Figure S6.**

### Figure S7

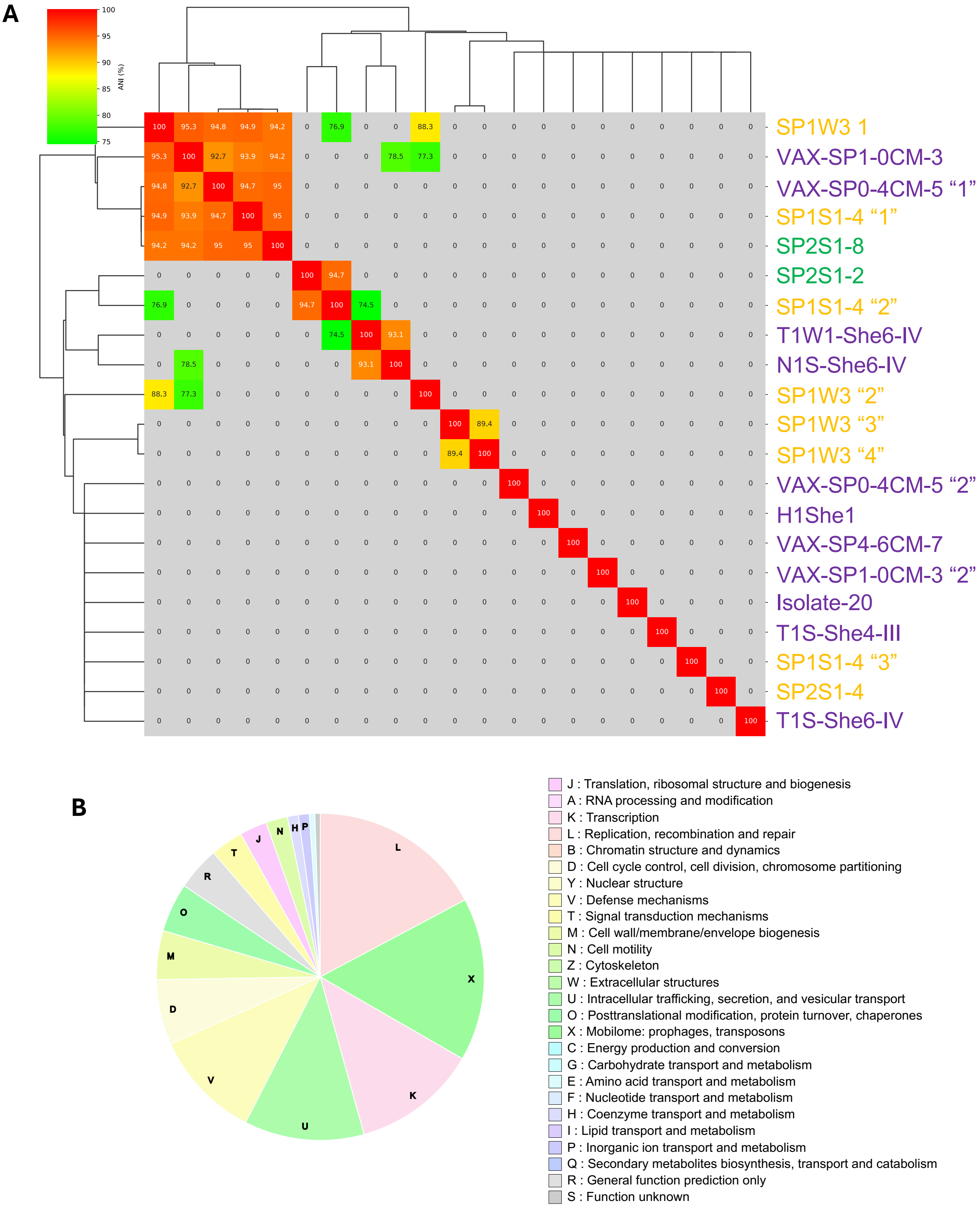

### Figure S8

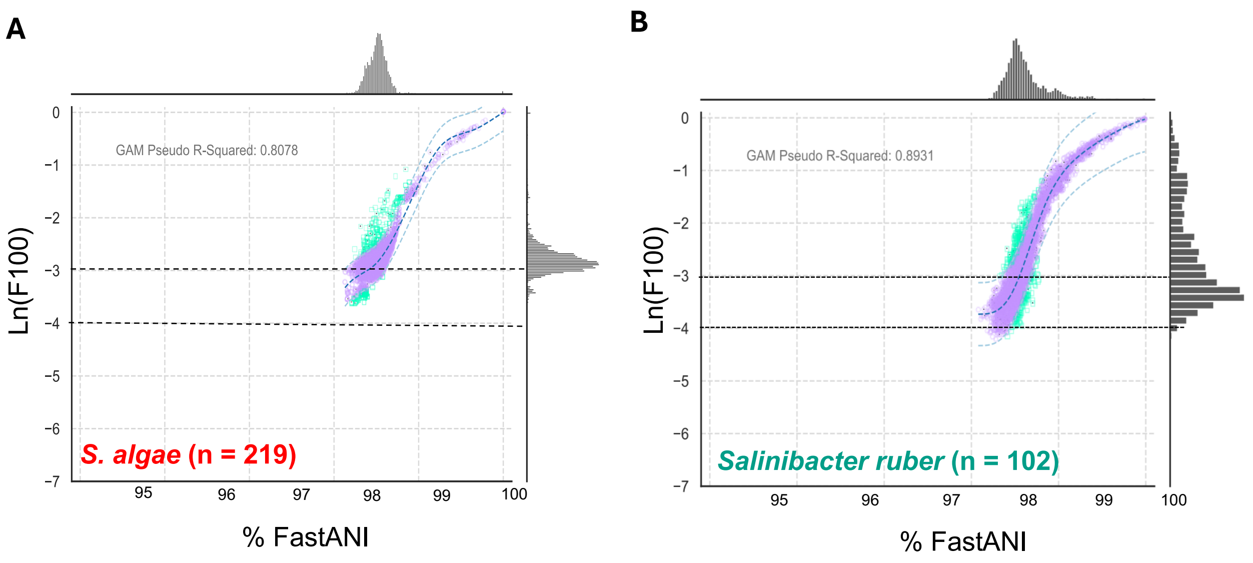

### Figure S9

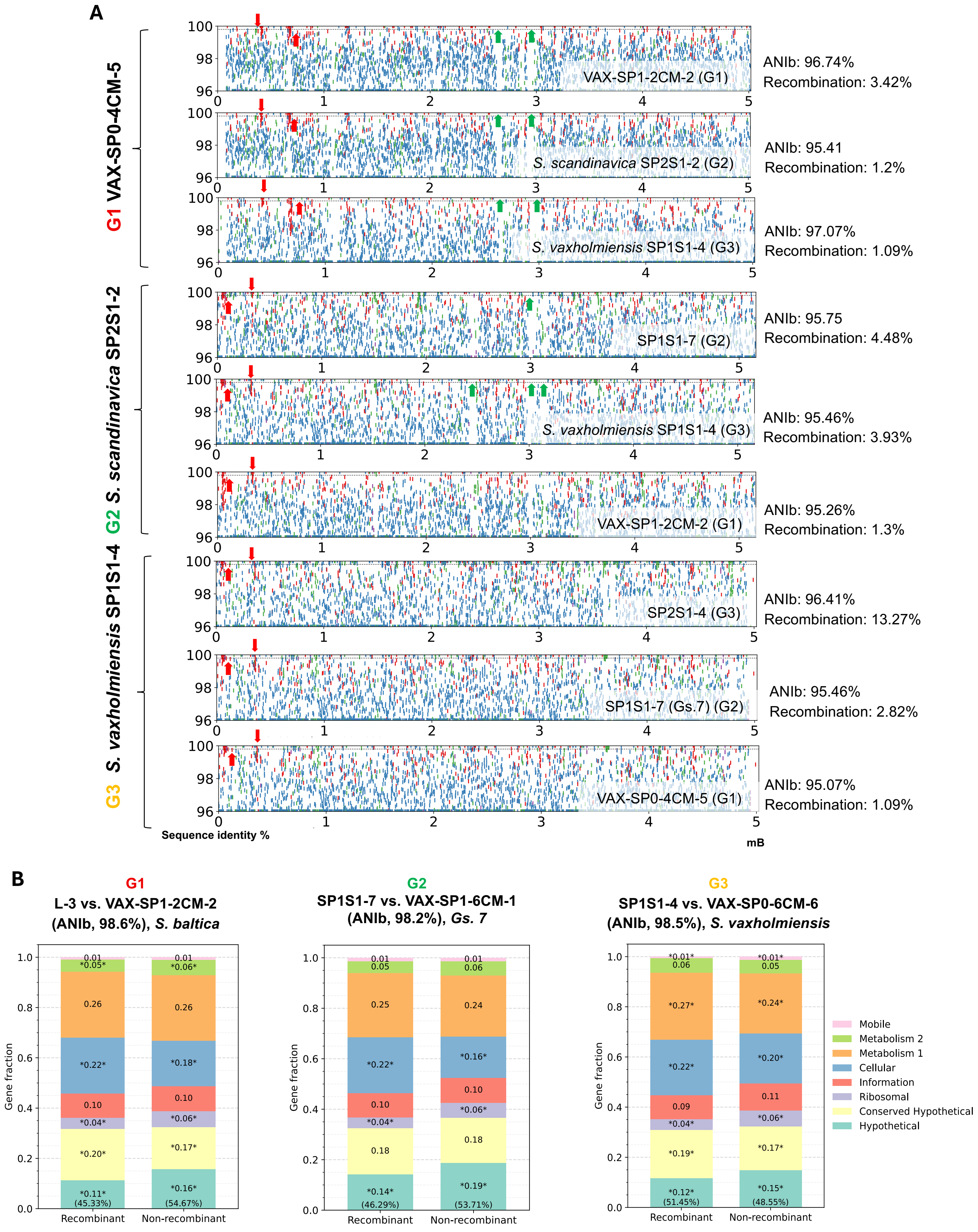

### Figure S10

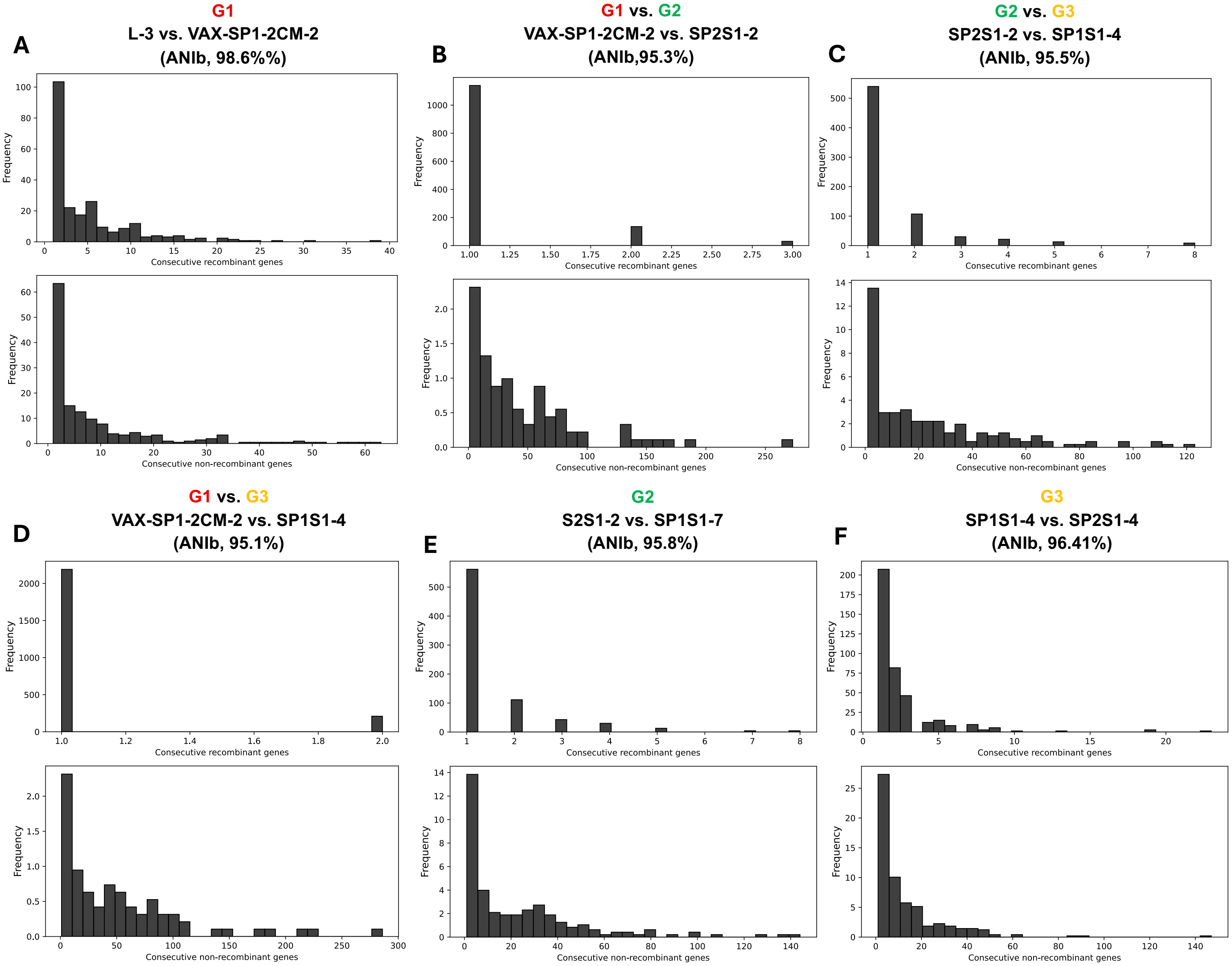
